# Thinking ahead: spontaneous prediction in context as a keystone of language in humans and machines

**DOI:** 10.1101/2020.12.02.403477

**Authors:** Ariel Goldstein, Zaid Zada, Eliav Buchnik, Mariano Schain, Amy Price, Bobbi Aubrey, Samuel A. Nastase, Amir Feder, Dotan Emanuel, Alon Cohen, Aren Jansen, Harshvardhan Gazula, Gina Choe, Aditi Rao, Se Catherine Kim, Colton Casto, Lora Fanda, Werner Doyle, Daniel Friedman, Patricia Dugan, Lucia Melloni, Roi Reichart, Sasha Devore, Adeen Flinker, Liat Hasenfratz, Omer Levy, Avinatan Hassidim, Michael Brenner, Yossi Matias, Kenneth A. Norman, Orrin Devinsky, Uri Hasson

## Abstract

Departing from traditional linguistic models, advances in deep learning have resulted in a new type of predictive (autoregressive) deep language models (DLMs). Using a self-supervised next-word prediction task, these models are trained to generate appropriate linguistic responses in a given context. We provide empirical evidence that the human brain and autoregressive DLMs share three fundamental computational principles as they process natural language: 1) both are engaged in continuous next-word prediction before word-onset; 2) both match their pre-onset predictions to the incoming word to calculate post-onset surprise (i.e., prediction error signals); 3) both represent words as a function of the previous context. In support of these three principles, our findings indicate that: a) the neural activity before word-onset contains context-dependent predictive information about forthcoming words, even hundreds of milliseconds before the words are perceived; b) the neural activity after word-onset reflects the surprise level and prediction error; and c) autoregressive DLM contextual embeddings capture the neural representation of context-specific word meaning better than arbitrary or static semantic embeddings. Together, our findings suggest that autoregressive DLMs provide a novel and biologically feasible computational framework for studying the neural basis of language.

## Introduction

The outstanding success of autoregressive (predictive) deep language models (DLMs) is striking from a theoretical and practical perspective because they have emerged from a very different scientific paradigm than traditional psycholinguist models^1^. In traditional psycholinguistic approaches, human language is explained with interpretable models that combine symbolic elements (e.g., nouns, verbs, adjectives, adverbs, etc.) with rule-based operations ^2, 3^. In contrast, autoregressive DLMs learn language from real-world textual examples “in the wild,” with minimal or no explicit prior knowledge about language structure. Autoregressive DLMs do not parse words into parts of speech or apply explicit syntactic transformations. Rather, they learn to encode a sequence of words into a numerical vector, termed a contextual embedding, from which the model decodes the next word. After learning, the next-word prediction principle allows the generation of well-formed texts in entirely new contexts never seen during training ^1, 5, 6^.

Autoregressive DLMs have proven to be extremely effective in capturing the structure of language ^4, 7–9^. It is unclear, however, if the core computational principles of autoregressive DLMs are related to the way the human brain processes language. Past research has leveraged language models and machine learning to extract semantic representation in the brain ^10–18^. But such studies did not view autoregressive DLMs as feasible cognitive models for how the human brain codes language. In contrast, recent theoretical papers argue that there are fundamental connections between DLMs and how the brain processes language ^1, 19, 20^.

In agreement with such a theoretical perspective, we provide novel empirical evidence that the human brain processes incoming speech similar to an autoregressive DLM (Fig. 1). In particular, the human brain and autoregressive DLMs share three computational principles: I) both are engaged in continuous context-dependent next-word prediction before word-onset; II) both match pre-onset predictions to the incoming word to induce post-onset surprise (i.e., prediction-error signals); III) both represent words using contextual embeddings. The main contribution of this study is the provision of direct evidence for the continuous engagement of the brain in next-word prediction before word-onset (computational principle I). In agreement with recent publications ^14, 16, 21–24^, we also provide novel neural evidence in support of computational principles II and III.

**Figure 1.**
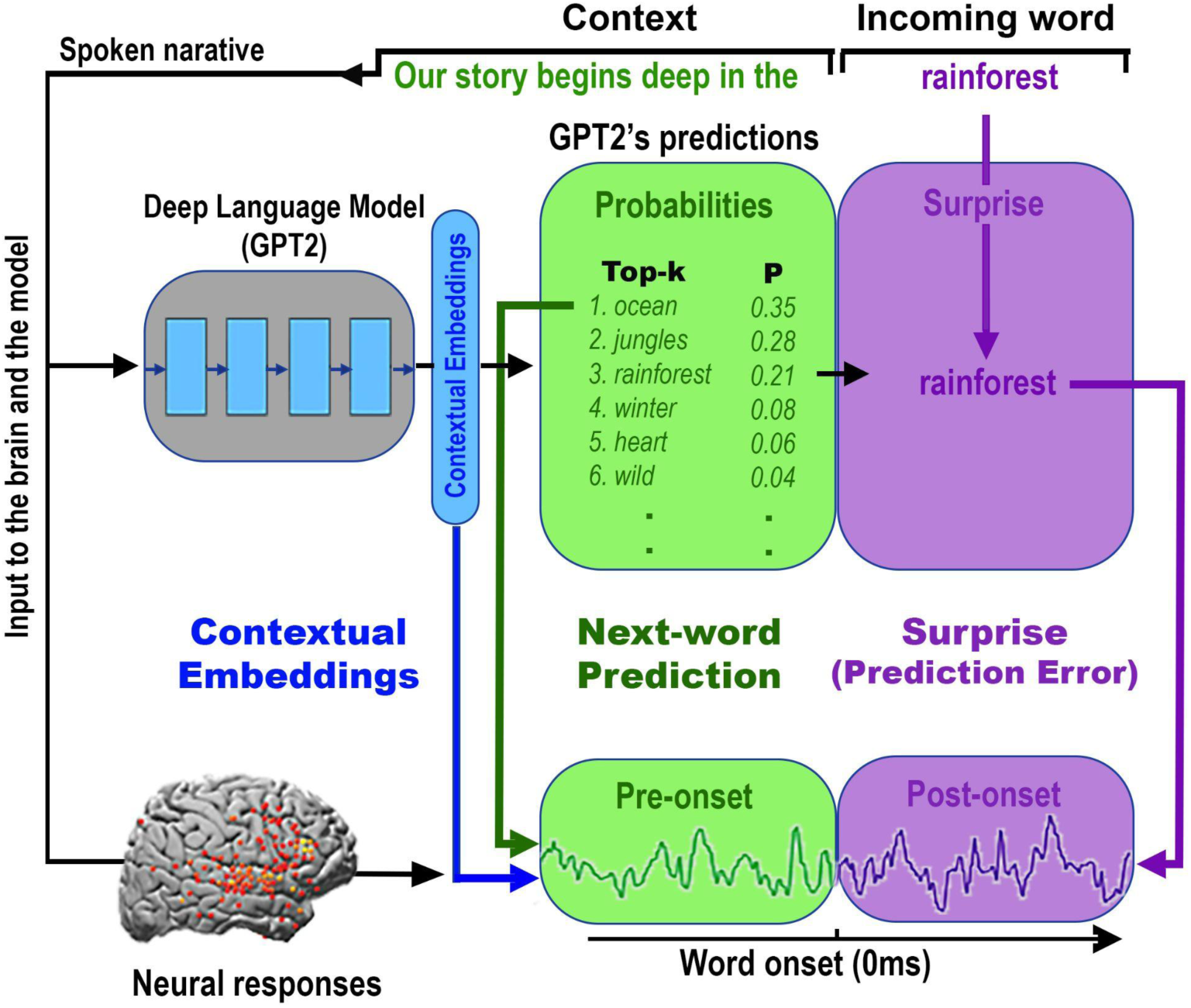
Shared computational principles between the brain and autoregressive deep language models (DLMs) in processing natural language. For each sequence of words in the text, GPT-2 generates a contextual embedding (blue), which is used to assign probabilities to the identity of the next word (green box). Next, GPT-2 uses its pre-onset predictions to calculate its surprise level (i.e., error signal) when the next word is revealed (purple box). The minimization of the surprise is the mechanism by which GPT-2 is trained to generate well-formed outputs. Each colored box and arrow represents the relationship between a given computational principle of the autoregressive DLM and the neural responses. The green boxes represent that both the brain and autoregressive DLMs are engaged in context-dependent prediction of the upcoming word before word onset. The green arrow indicates that we take the actual predictions from GPT-2 (e.g. the top-1 prediction ”ocean” for the example sentence “our story begins deep in the…”) to model the neural responses before word onset (Fig. 4). The purple boxes represent that both the brain and autoregressive DLMs are engaged in the assessment of their predictions after word onset. The purple arrow indicates that we take the actual perceived next word (“forest” in our example), as well as GPT-2’s surprise level for the perceived word (cross-entropy) to model the neural responses after word onset (Fig. 4B and 5B). The blue boxes represent that both the brain and autoregressive DLMs use contextual embeddings to represent words. The blue arrow indicates that we take the contextual embeddings from GPT-2 to model the neural responses (Fig. 6 & 8). We argue here that, although the internal word-processing mechanisms are not the same for the brain and DLMs, they do share three core computational principles: 1) continuous context-dependent next-word prediction before word-onset; 2) reliance on the pre-onset prediction to calculate post-word-onset surprise; and finally, 3) context-specific representation of meaning.

### Computational Principle I: Like autoregressive DLMs, neural activity <u>before word onset</u> reflects context-dependent next-word prediction (Fig. 1, green)

The constant engagement in next-word prediction before word-onset is the bedrock objective of autoregressive DLMs ^4, 7–9, 25^. Similarly, the claim that the brain is constantly engaged in predicting the incoming input is fundamental to numerous predictive coding theories ^26–30^. However, even after decades of research, behavioral and neural evidence for the brain’s propensity to predict upcoming words before they are perceived has remained indirect. On the behavioral level, the ability to predict upcoming words has been mostly tested with highly-controlled sentence stimuli (i.e., the cloze procedure ^31–35^). Thus, we still do not know how accurate listeners’ predictions are in open-ended natural contexts. The first section of the paper provides novel behavioral evidence that humans can predict forthcoming words in a natural context with remarkable accuracy, and that, given a sufficient context window, next-word predictions in humans and an autoregressive DLM (GPT-2^4^) match. On the neuronal level, we provide novel evidence that the brain is spontaneously engaged in next-word prediction before word-onset. In addition, we were able to track the content of the predictions even when they were inaccurate. These findings provide the missing evidence that the brain, like autoregressive DLMs, is constantly involved in next-word prediction before word-onset as it processes natural language.

### Computational Principle II: Like autoregressive DLMs, the brain relies on pre-onset predictions to calculate post-word-onset surprise (prediction-error) signals (Fig. 1, purple)

Detecting increased neural activity 400 ms after word-onset for unpredictable words, documented across many studies ^36–40^, has traditionally been used as indirect evidence for pre-onset predictions. Recent development of autoregressive DLMs, like GPT-2, provides a powerful new way to quantify the surprise and confidence levels for each upcoming word in natural language ^14, 22, 23^. Specifically, autoregressive DLMs, like GPT-2, provide a unified framework that connects pre-onset next-word prediction with post-onset prediction-error and surprise. Autoregressive DLMs use the confidence in pre-onset next-word predictions to calculate post-onset surprise level. If the incoming word received a high probability before word onset, then the resulting surprise (i.e., prediction error ) would be low, and vice versa for prediction with low probabilities. The terms “surprise” and “prediction error” are closely related. In accordance with prior literature^46^, we will use the term surprise throughout the paper. For example, in Fig. 1, the surprise level assigned to the incoming word “rainforest” (Fig. 1, purple box) is determined by the assigned probability to that word before word onset (Fig. 1, green box). Here we used the spatiotemporally-resolved intracranial recordings to not only replicate the previous findings ^14, 22, 23^, but also to map the temporal coupling between confidence in pre-onset prediction and post word-onset surprise signals. Further, we demonstrate enhanced encoding (i.e., higher correlations between neural signals and embeddings) for unpredictable words in higher-order language areas 400 ms post-word-onset. Together, these findings provide compelling evidence that, similar to DLMs, the biological neural error signals after word onset are coupled to pre-onset neural signals associated with next-word predictions.

### Computational Principle III: Like autoregressive DLMs, the brain represents words contextually (Fig. 1, blue)

Autoregressive DLMs encode the unique, context-specific meaning of words based on the sequence of prior words. Concurrent findings demonstrate that contextual embeddings derived from GPT-2 provide better modeling of neural responses in multiple brain areas than static word embeddings (i.e., non-contextual) ^16, 17^. Our paper goes beyond these findings by showing that contextual embeddings encapsulate information about past contexts as well as next-word predictions. Dissociating context and next-word prediction allows us to use contextual embeddings both to model pre-onset neural activity associated with context representation, and to model pre-onset neural activity associated with next-word predictions. Finally, we demonstrate that contextual embeddings are better than non-contextual embeddings at predicting word identity from cortical activity (i.e., decoding) before (and after) word onset.

Taken together, our findings provide compelling evidence for core computational principles of pre-onset prediction, post-onset surprise, and contextual representation, shared by autoregressive DLMs and the human brain. These results support a new unified modeling framework for the study of the neural basis of the human language faculty.

## Results

### Section I - Prediction before word-onset

#### Comparison of next-word prediction behavior in autoregressive DLMs and humans

We developed a novel sliding-window behavioral protocol to directly quantify humans’ ability to predict every word in a natural context (Fig. 2A-B). 50 participants proceeded word-by-word through a 30-minute transcribed podcast (“Monkey in the Middle”, *this American Life* podcast^41^) and provided a prediction of each upcoming word. The procedure yielded 50 predictions for each of the story’s 5113 words (see Fig. 2C, and Materials and Methods). We calculated the mean prediction performance for each word in the narrative, which we refer to as a “predictability score” (Fig. 2D). A predictability score of 100% indicates that all participants correctly guessed the next word and a predictability score of 0% indicates that no participant predicted the upcoming word. This allowed us to address the following questions: First, how good are humans at next-word prediction? Second, how much do human predictions align with autoregressive DLM predictions?

**Figure 2.**
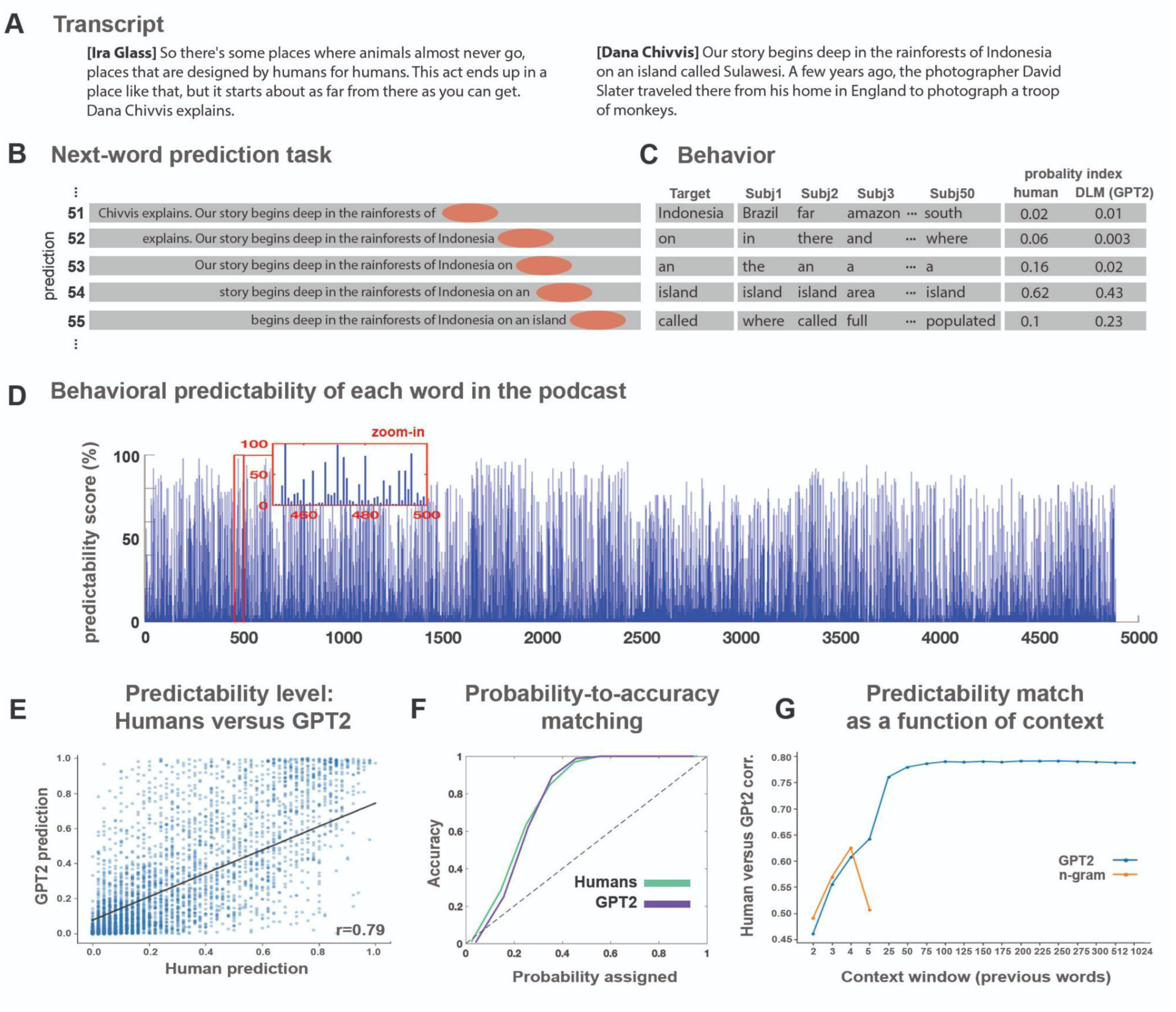
Behavioral assessment of the human ability to predict forthcoming words in a natural context. **A)** The stimulus was transcribed for the behavioral experiment. **B)** A 10-word sliding window was presented in each trial, and participants were asked to type their prediction of the next word. Once entered, the correct word is presented, and the window slides forward by one word. **C)** For each word, we calculated the proportion of participants that predicted the forthcoming word correctly. **D)** Human’s predictability scores across words. **E)** Human’s predictability scores versus GPT-2’s predictability scores for each upcoming word in the podcast. **F)** Match between humans’ (green) and GPT-2’s (purple) assigned probability and the actual accuracy for their top-1 predictions. **G)** Correlation between human predictions and GPT-2 predictions (as reported in panel D) for different context window lengths ranging from 2–1024 preceding tokens (blue line). Correlation between human predictions and 2- to 5-gram model predictions (red line).

#### Word-by-word behavioral prediction during a natural story

Participants were able to predict many upcoming words in a complex and unfamiliar story. The average human predictability score was 28%, SE = 0.5%, in comparison to a predictability score of 6% for blindly guessing the most frequent word in the text (“the”). About 600 words had a predictability score higher than 70%. Interestingly, high predictability was not confined to the last words in a sentence and applied to words from all parts of speech (21.44% nouns, 14.64% verbs, 41.62% function words, 4.35% adjectives, adverbs, and 17.94% other). This suggests that humans are proficient in predicting upcoming words in real-life contexts when asked to do so.

#### Comparing human and DLM next-word predictions and probabilities

Autoregressive DLMs learn how to generate well-formed linguistic outputs by improving their ability to predict the next word in natural linguistic contexts. We compared human and autoregressive DLM predictions of the words of the podcast as a function of prior context. As an instance of an autoregressive DLM, we chose to work with GPT-2^4^. GPT-2 is a pre-trained autoregressive language model with state-of-the-art performance on tasks related to reading comprehension, translation, text summarization, and question answering. GPT-2 is trained by maximizing the log-probability of a token given its 1024 past tokens (for full description see^8^). For each word in the transcript, we extracted the most probable next-word prediction and the prediction probability of the actual word assigned by GPT-2 as a function of context (i.e., maximum context window of 1024 tokens). For example, GPT-2 assigned a probability of 0.82 to the upcoming word “*monkeys*” when it received the preceding words in the story as contextual input: “… So after two days of these near misses, he changed strategies. He put his camera on a tripod and threw down some cookies to try to entice the .” Human predictability scores and GPT-2 estimations of predictability were highly correlated (Fig. 2E, *r* = .79, *p* < .001). In this case, both GPT-2’s and human’s most probable next-word prediction was “monkeys”. In 49.1% of the cases the human’s most probable prediction and GPT-2’s most probable prediction matched (irrespective of accuracy). For baseline comparison we report the same agreement measure with human prediction for 2- to 5-gram models in Fig. S1 (see Materials and Methods). In terms of accuracy, GPT-2 and humans jointly predicted correctly 27.6% of the words and jointly predicted incorrectly 54.7% of the words. Only 9.2% of the words humans predicted correctly were not correctly predicted by GPT-2, and only 8.4% of the words correctly predicted by GPT-2 were not correctly predicted by humans (see Fig. S2).

Finally, we compared the match between the confidence level and the accuracy level of GPT- 2’s and humans’ predictions. For example, if the model (or humans) assigned a 20% probability, would it (or they) produce only 20% correct predictions? Both humans and GPT-2 had a remarkably similar confidence-to-accuracy function (Fig. 2F). Specifically, GPT-2 and humans were under-confident in their predictions and above 95% correct when the probabilities were higher than 40%. These analyses suggest that GPT-2’s and humans’ next-word predictions are similar in natural contexts.

#### Prediction as a function of contextual window size

In natural comprehension (e.g., listening to or reading a story), predictions for upcoming words are influenced by information accumulated over multiple timescales: from the most recent words to the information gathered over multiple paragraphs ^42^. We tested if GPT-2’s predictions would improve as a function of the context window as it does in humans. To that end, we varied GPT-2’s input window size (from two tokens up to 1024 tokens) and examined the impact of contextual window-size on the match with human behavior. The correlation between human and GPT-2 word predictions improved as the contextual window increased (from *r* = .46, *p* < .001 at two-word context to an asymptote of *r* = .79 at 100-word context; Fig. 2G). For baseline comparison we also plot the correlations of 2- to 5-gram models with human predictions (Fig. 2G; also see Materials and Methods).

#### Neural activity before word onset reflects next-word predictions

The behavioral study indicates that listeners can accurately predict upcoming words in a natural open-ended context when explicitly instructed. Furthermore, it suggests human predictions and autoregressive DLM predictions are matched in natural contexts. Next, we asked whether the human brain, like autoregressive DLM, is continuously engaged in spontaneous next-word prediction before word onset without explicit instruction. To that end, we used electrocorticography (ECoG) signals from nine epileptic patients who volunteered to participate in the study (see Fig. 3A for a map of all electrodes). All participants listened to the same spoken story used in the behavioral experiment. In contrast to the behavioral study, the patients engaged in free listening—with no explicit instructions to predict upcoming words. Comprehension was verified using a post-listening questionnaire. Across participants, we had better coverage in the left hemisphere (1106 electrodes) than in the right hemisphere (233 electrodes). Thus, we focus on language processing in the left hemisphere, but we also present the encoding results for the right hemisphere in the supplementary materials (Fig. S3). The raw signal is preprocessed to reflect high-frequency broadband (70–200Hz) power activity (for full preprocessing procedures see Materials and Methods).

**Figure 3.**
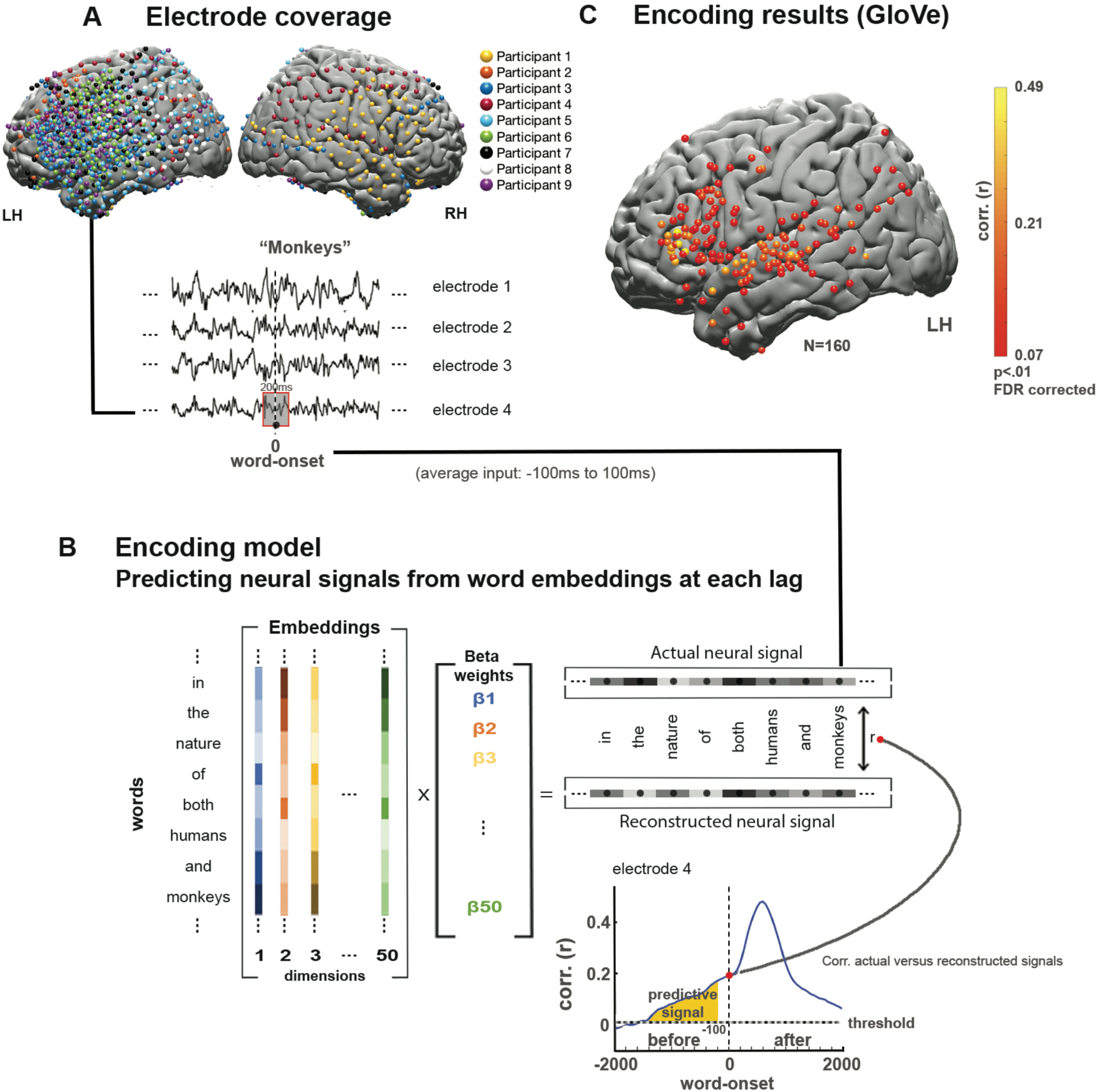
Linear encoding model used to predict the neural responses to each word in the narrative before and after word-onset. **A)** Brain coverage consisted of 1339 electrodes (across nine participants). The words are aligned with the neural signal; each word’s onset (moment of articulation) is designated at lag 0. Responses are averaged over a window of 200 ms and provided as input to the encoding model. **B)** A series of 50 coefficients corresponding to the features of the word embeddings is learned using linear regression to predict the neural signal across words from the assigned embeddings. The model was evaluated by computing the correlation between the reconstructed signal and the actual signal for a held out test word. This procedure was repeated for each lag and each electrode, using a 25 ms sliding window. The dashed horizontal line indicates the statistical threshold (q < .01 FDR corrected). Lags of -100 ms or more preceding word onset contain only neural information sampled before the word was perceived (yellow color). **C)** Electrodes with significant correlation at the peaked lag between predicted and actual word responses for semantic embeddings (GloVe).

Below we provide multiple lines of evidence that the brain, like autoregressive DLMs, is spontaneously engaged in next-word prediction before word onset. The first section focuses solely on the pre-onset prediction of individual words by using static (i.e., non-contextual) word embeddings (GloVe^43^ and word2vec^44^). In the third section, we investigate how the brain adjusts its responses to individual words as a function of context, by relying on contextual embeddings.

#### Localizing neural responses to natural speech using static embeddings

We used a linear encoding model and static semantic embeddings (GloVe) to localize electrodes containing reliable responses to single words in the narrative (see Fig. 3A-B and Materials and Methods). Encoding models learn a mapping to predict brain signals from a representation of the task or stimulus^45^. The model identified 160 electrodes in the left hemisphere (LH) with significant correlations (after correction for multiple comparisons, see Materials and Methods for details). Electrodes were found in early auditory areas, motor cortex, and language areas (see Fig. 3C for LH electrodes, and Fig. S3 for right hemisphere (RH) electrodes).

#### Encoding neural responses before word onset

In the behavioral experiment (Fig. 2), we demonstrated people’s capacity to predict upcoming words in the story. Next, we tested whether the neural signals also contain information about the identity of the predicted words before they are perceived (i.e., before word-onset). The word-level encoding model (based on GloVe word embeddings) yielded significant correlations with predicted neural responses to upcoming words up to 800 ms before word-onset (Fig. 4A; for single electrodes encoding models see Fig. S4). Peak encoding performance was observed 150–200 ms after word onset (lag 0), but the models performed above chance up to 800 ms before word-onset. As a baseline for the noise level, we randomly shuffled the GloVe embeddings, assigning a different vector to the occurrence of each word in the podcast. The analysis yields a flat encoding around zero (Fig. 4A). The encoding results using GloVe embeddings were replicated using 100-dimensional static embeddings from word2vec (Fig. S5). To control for the contextual dependencies between adjacent words in the GloVe embeddings, we demonstrate that the significant encoding before word onset holds even after removing the information of previous GloVe embedding (Fig. S6-A). This supports the claim that the brain continuously predicts semantic information about upcoming words’ meaning before they are perceived.

**Figure 4.**
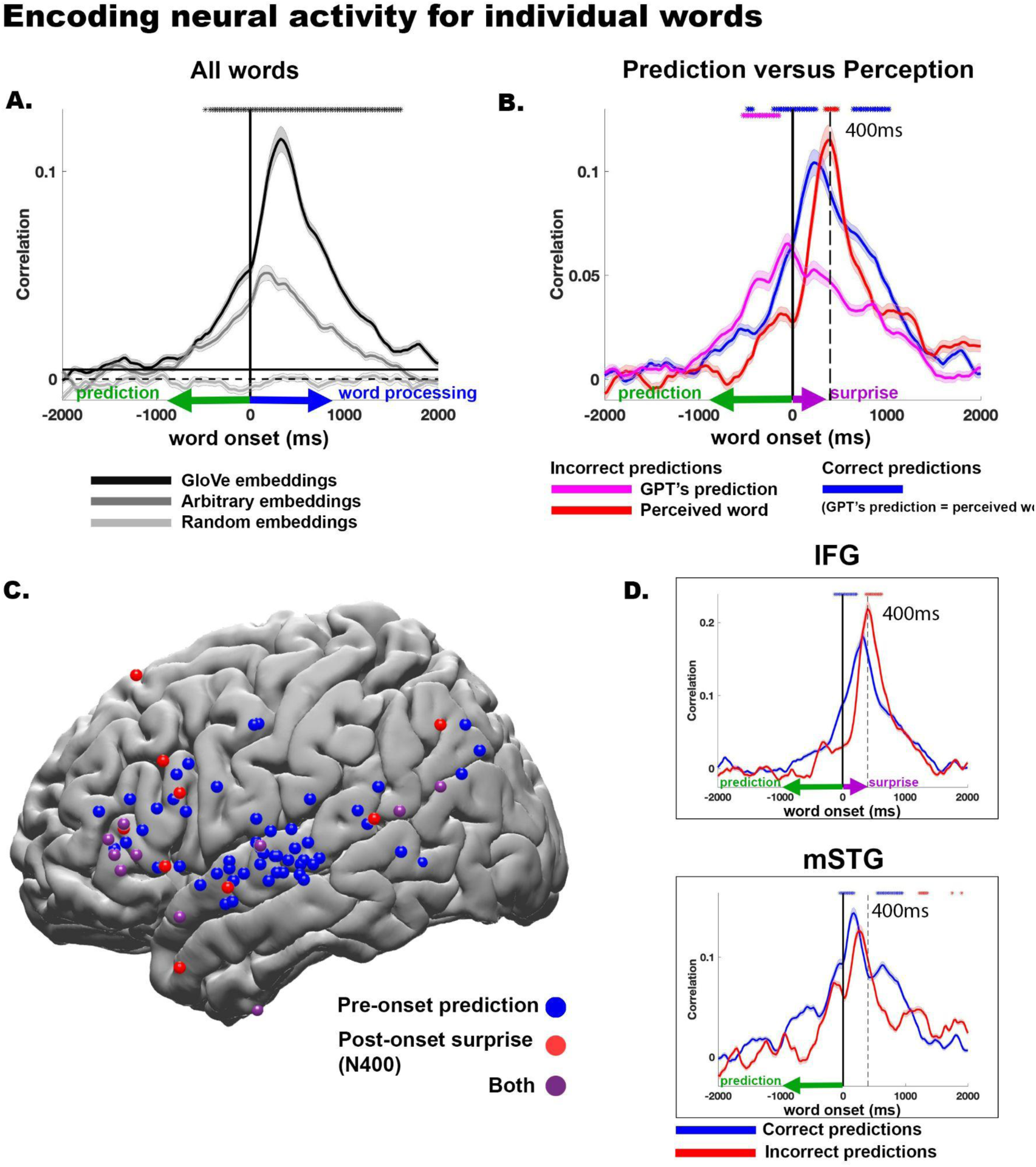
Modeling of neural signals before and after word onset for predictable, unpredictable, and incorrectly predicted words. **A)** Estimating neural signals for all individual words from word embeddings (encoding). The encoding analysis was performed in each electrode with significant encoding for GloVe embeddings (n=160), and then averaged across electrodes (see map of electrodes in Fig. 3c). The error bars indicate the standard error of the encoding models across electrodes. Using arbitrary embeddings we managed to encode information as to the identity of the incoming word before and after word-onset. Using word embeddings (GloVe), which contain contextual information as to the relation among words in natural language, further improves the encoding models before and after word-onset. Furthermore, we observed a robust encoding to upcoming words starting -1000 ms before word onset. The horizontal continuous black line specifies the statistical threshold. Black asterisks indicate lags for which the encoding based on GloVe embeddings significantly outperforms the encoding based on arbitrary embeddings. **B)** Estimating neural signals for correctly predicted words (blue), incorrectly predicted words (magenta), and the actual unexpected perceived word (red). Note that encoding before word onset was aligned with the content of the predicted words, whereas the encoding after word onset was aligned with the content of the perceived words. Moreover, we observed an increase in encoding performance for surprising words compared to predicted words 400 ms after word onset. Magenta asterisks represent significant differences between incorrect GPT-2 predictions (magenta line) and correct predictions (blue line). Red asterisks represent significantly higher values for incorrectly predicted words (red line) than correctly predicted words (blue line). Blue asterisks represent significantly higher values for correctly predicted words (blue line) than incorrectly predicted (red line). **C)** Single electrodes which show significant pre-onset prediction for correct words are marked in blue (pre-onset prediction). Single electrodes that show a significant increase in encoding for incorrect words over correct words 400 ms after word onset are marked in red (post-onset surprise). Electrodes that show both pre-onset prediction signals and post-onset surprise signals are marked in purple (both). **D)** Average estimation of activity for all electrodes in the inferior frontal gyrus (IFG) and the middle superior temporal gyrus (mSTG).

To test whether GloVe based encoding is affected by the semantic knowledge embedded in the model we shuffled the word embeddings. Interestingly, when assigning a non-match GloVe embedding (from the story) to each word such that multiple occurrences of the same word received the same (but non-match) GloVe embedding the encoding decreases (Fig. S7). This indicates that the relational linguistic information encoded in GloVe embeddings is also embedded in the neural activity.

#### Encoding neural responses before word onset independent of context

To test if the significant encoding before word onset is driven by contextual dependencies between adjacent words in the GloVe embeddings, we also trained encoding models to predict neural responses using 50-dimensional static arbitrary embeddings, randomly sampled from a uniform [-1,1] distribution. Arbitrary embeddings effectively remove the contextual information about the statistical relationship between words included in GloVe embeddings (Fig. 4A). Even for arbitrary embeddings, we were able to obtain significant encoding before word onset as to the identity of the upcoming word (for single electrodes encoding models see Fig. S4). To make sure that the analysis does not rely on local dependencies among adjacent words, we repeated the arbitrary based encoding analysis after removing the first word from any bi-gram that repeats more than once in the dataset (Fig. S6-B). The ability to encode the neural activity for the upcoming words before word onset with the arbitrary embeddings remained significant.

To further demonstrate that predicting the next word before word onset goes above and beyond the contextual information embedded in the previous word, we ran an additional control analysis. In the control analysis, we encode the neural activity using the arbitrary word embedding assigned to the previous word (Fig. S6-C, blue). Next, we ran an encoding model using the concatenation of the previous and current word embeddings (Fig. S6-C, red). We found a significant difference between these two models before word onset. This indicates that the neural responses before word-onset contain information related to the next word above and beyond the contextual information embedded in the previous word. Together, these results suggest that the brain is constantly engaged in the prediction of upcoming words before they are perceived as it processes natural language.

#### Neural signals before word onset contain information about listeners’ incorrect predictions

Pre-onset activity associated with next-word prediction should match the prediction content even when the prediction turned out to be incorrect. In contrast, post-onset activity should match the content of the incoming word, even if it was unpredicted. To test this hypothesis we divided the signal into correct and incorrect predictions using GPT-2 (see Materials and methods) and computed encoding models. We also ran the same analyses using human predictions. We modeled the neural activity using: 1) the GloVe embeddings of the correctly predicted words (blue in Fig. 4B). In this condition the pre-onset word prediction matches the identity of the perceived incoming word; 2) the GloVe embedding for the incorrectly predicted words (magenta in Fig. 4B); 3) Since in the incorrect predictions condition the predicted word does not match the identity of the perceived word, we also modeled the signal using the GloVe embedding of the actual unpredictable words humans perceived (red in Fig. 4B).

The neural responses before word onset contained information about human predictions as to the identity of the next word. Crucially, the encoding was high for both correct and incorrect predictions (Fig. 4B, S8 blue and magenta). This demonstrates that pre-word-onset neural activity contains information about what listeners actually predicted, irrespective of what they subsequently perceived. Similar results were obtained using human predictions (Fig. S8). In contrast, the neural responses after word onset contained information about the words that were actually perceived, irrespective of GPT-2’s predictions (Fig. 4B, blue and red). The analysis of the incorrect predictions unequivocally disentangles the pre-word onset processes associated with word prediction from the post-word onset comprehension-related processes. It demonstrates that neural signals before word-onset contain information about listeners’ internal expectations. Furthermore, it demonstrates how autoregressive DLMs’ predictions can be used for modeling humans’ predictions at the behavioral and neural levels.

In summary, these multiple pieces of evidence, which are based on encoding analyses, suggest that the brain, like autoregressive DLM, is constantly predicting the next word before its onset as it processes incoming natural speech. Next, we provide more evidence for coupling between pre-onset prediction and post-onset surprise level and error signals.

Electrodes were selected based on anatomical location irrespective of their selectivity. Note the strong coupling between pre-onset prediction and post-onset surprise along the IFG, but not the mSTG, as indicated by the significant difference between the unexpected and predicted words (red asterisks) around 400 ms post onset. The comparison was done for each leg separately and results were FDR corrected (q < .01) (See Materials and Methods).

### Section II - Connecting pre-onset predictions with post-onset surprise signals

Autoregressive language models provide a unified framework for modeling pre-onset next-word predictions and post-onset surprise (i.e., prediction error signals). We used pre-trained GPT-2’s internal estimates for each upcoming word (Fig. 1, green) to establish a connection between pre-onset prediction and post-onset surprise at the neural level.

#### Increased semantic processing for unpredictable words 400 ms after word onset along the IFG

We started by demonstrating enhanced processing 400 ms after word onset for surprising words that GPT-2 failed to predict (Fig. 4B, red versus blue lines). This enhanced neural response to unpredictable words after word-onset may reflect the calculations of error signal. Next, we asked whether the increase of post-onset processing for surprising words (i.e., higher values in the encoding model) would be localized to higher-order language areas. To that end we mapped single electrodes that show: 1) pre-onset encoding for upcoming words (i.e., prediction effect, Fig. 4C blue); 2) post-onset increase encoding for incorrect predictions over correct predictions 400 ms after word onset (i.e., surprise, Fig. 4C red); 3) encode both signals associated with pre-onset prediction and post-onset prediction error (Fig. 4C purple). The map shows a widespread prediction effect across all language areas, specifically, middle superior temporal gyrus (mSTG; 28 electrodes) with an additional increase in processing of incorrect words along the IFG (23 electrodes) (Fig. 4C-D). We calculated a two-way ANOVA for the signals at 400 ms lag. We defined a within-electrode factor of accuracy (correct/incorrect), and a between-electrode factor of brain region (mSTG/IFG). We found a significant two-way interaction (F(1,49) = 8.42, p = .006), suggesting a larger difference between correct and incorrect predictions in IFG than in mSTG at 400 ms lag.

#### Increased activity as a function of surprise level 400 ms after word onset

Autoregressive DLMs, such as GPT-2, use their pre-onset predictions to calculate the post-onset surprise level as to the identity of the incoming word. It was already shown that the activation level post-onset is correlated with the surprise level ^14, 21–23, 46^. We replicated this finding in our data. In addition, our high-quality intracranial recordings allowed us to link pre-onset confidence level and the post-onset surprise level. Pre-onset confidence level was assessed using entropy (see Materials and Methods), which is a measure of GPT-2’s uncertainty] level in its prediction before word-onset. High entropy indicates that the model is uncertain about its predictions, whereas low entropy indicates that the model is confident. Post-onset surprise level was assessed using a cross-entropy measure that depends on the probability assigned to the incoming word before it is perceived (see examples in Fig. 1, purple box and Materials and Methods). Assigning a low probability to the word before word onset will result in a post-onset high surprise when the word is perceived, and vice versa for high probability words.

Pre-onset activity (using the same 160 electrodes used for Fig. 4A-B) increased for correct predictions, whereas, in agreement with prior research, post-onset activity increased for incorrect predictions (Fig. 5A). The activity level was averaged for all words that were correctly (blue) or incorrectly (red) predicted. We observed increased activity for incorrect prediction 400 ms after word onset (Fig. 5A). In addition, GPT-2’s uncertainty (entropy) was negatively correlated with the activation level before word-onset (Fig. 5B, green). In other words, before onset, the higher the confidence (low uncertainty), the higher the activation level. In contrast, after word onset, the level of surprise (cross-entropy) was correlated with activation, and peaked around 400 ms (Fig. 5B, green). Since uncertainty correlates with surprise, we computed partial correlations between entropy, surprise, and neural signals. This analysis directly connects GPT-2’s internal predictions and neural activity before word onset and GPT-2’s internal surprise and the surprise (i.e., prediction error) embedded in the neural responses after word-onset.

**Figure 5.**
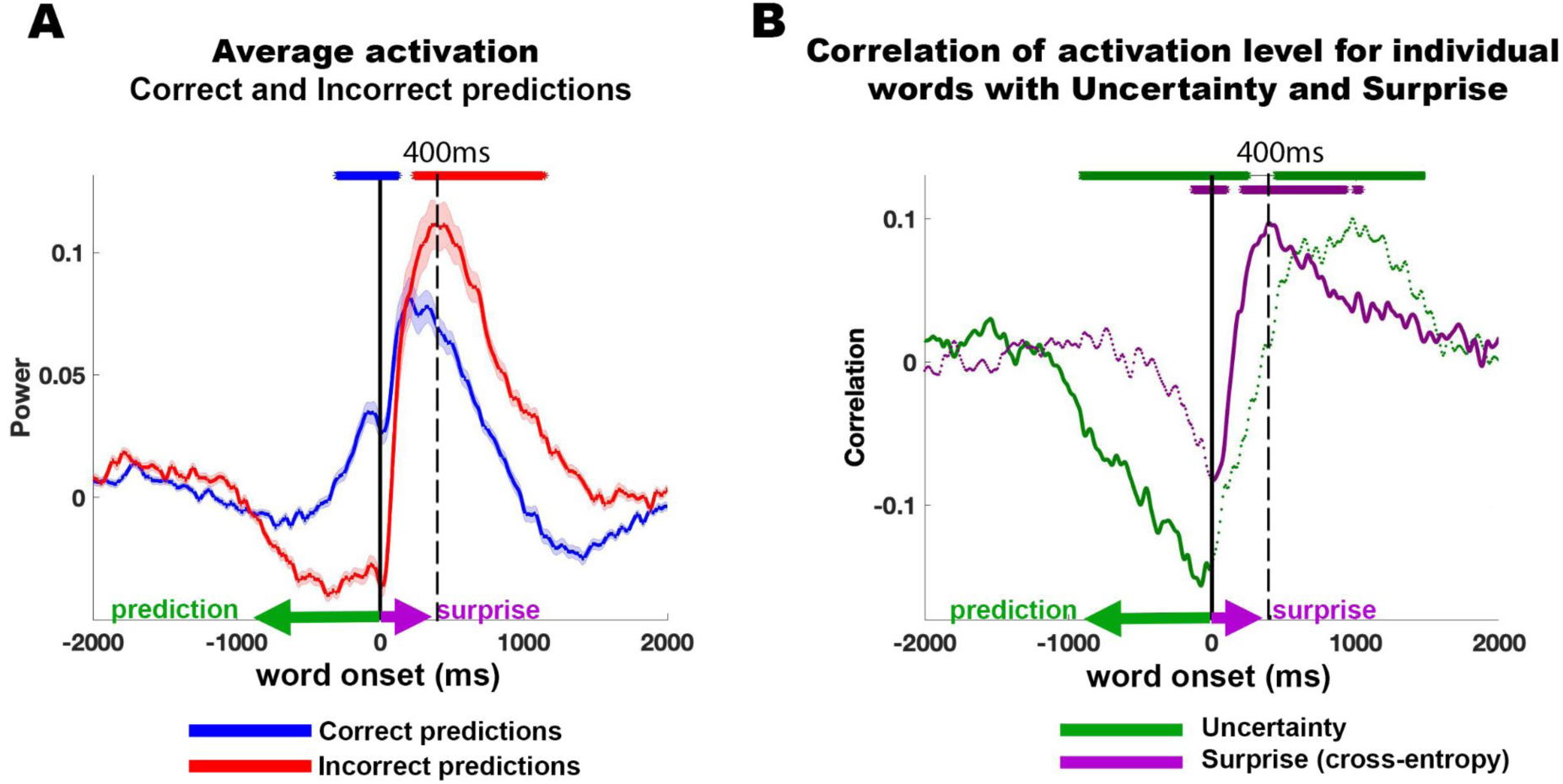
Uncertainty and surprise levels computed by GPT-2 correlate with pre-onset and post-onset neural activity respectively. **A)** Trigger-averaged activity (and standard error) across 160 electrodes with significant GloVe encoding for words (see Fig. 3C). Averaging was performed separately for words correctly predicted (blue) and incorrectly predicted (red). Note, the increase in signal activity for predictable words before onset, and for unpredictable words 400 ms after word onset. **B)** Partial correlations between uncertainty (entropy) and signal power controlling for cross-entropy (green line). Partial correlations between surprise (cross-entropy) and neural signals controlling for correlation with entropy (red,purple). Asterisks indicate correlation significance (FDR corrected, q<.01).

In summary, based on encoding and event-related activity, we introduce multiple pieces of evidence to link pre-onset next-word prediction processes with post-onset surprise processes using GPT-2’s internal estimates. This section further supports the claim that autoregressive DLMs can serve a theoretical framework for language comprehension-related processes. Next, we provide more evidence that GPT-2 tracks human neural signals and specifically that it represents words in a contextual dependent fashion, similar to humans.

### Section III - Contextual representation

#### Using contextual embeddings to predict neural responses to natural speech

Next-word prediction objective enables autoregressive DLMs to compress a sequence of words into a contextual embedding from which the model decodes the next word. The present results have established that the brain, similar to autoregressive DLMs, is also engaged in spontaneous next-word prediction as it listens to natural speech. Given this shared computational principle, we investigated whether the brain, like autoregressive DLMs, compresses word sequences into contextual representation.

In natural language, each word receives its full meaning based on the preceding words ^47–49^. For instance, consider how the word “shot” can have very different meanings in different contexts, such as “taking a shot with the camera”, “drinking a shot at the bar” or “making the game-winning shot”. Static word embeddings, like GloVe, assign one unique vector to the word “shot” and, as such, cannot capture the context-specific meaning of the word. In contrast, contextual embeddings assign a different embedding (vector) to every word as a function of its preceding words. Here we tested whether autoregressive DLMs that compress context into contextual embeddings provide a better cognitive model for neural activity during linguistic processing than static embeddings. To test this, we extracted the contextual embeddings from an autoregressive DLM (GPT-2) for each word in the story. To extract the contextual embedding of a word, we provided the model with the preceding sequence of all prior words (up to 1024 tokens) in the podcast and extracted the activation of the top embedding layer (see Materials and Methods).

#### Localizing neural responses to natural speech using contextual embeddings

Replacing static embeddings (GloVe) with contextual embeddings (GPT-2) improved encoding model performance in predicting the neural responses to words (Fig. 6A, S3). Encoding based on contextual embeddings resulted in statistically significant correlations in 208 electrodes in LH (and 34 in RH). 71 of these electrodes were not significantly predicted by the static embeddings (GloVe). The additional electrodes revealed by contextual embedding were mainly located in high-order language areas with long processing timescales along the inferior frontal gyrus, temporal pole, posterior superior temporal gyrus, parietal lobe, and angular gyrus ^42^. In addition, there was a noticeable improvement in the contextual embedding-based encoding model in the primary and supplementary motor cortices. The improvement is seen both at the peak of the encoding model and in the model’s ability to predict neural responses to words up to four seconds before word-onset (for the 160 electrodes with significant GloVe encoding; Fig. 6B, S4, and S9). The improvement in the ability to predict neural signals to each word while relying on autoregressive DLM’s contextual embeddings was robust and apparent even at the single electrode level (see Fig. S4 for a selection of electrodes). These results agree with concurrent studies demonstrating that contextual embeddings model neural responses to words better than static semantic embeddings ^15, 16, 50, 51^. Next, we asked which aspects of the contextual embedding drive the improvement in modeling the neural activity.

**Figure 6.**
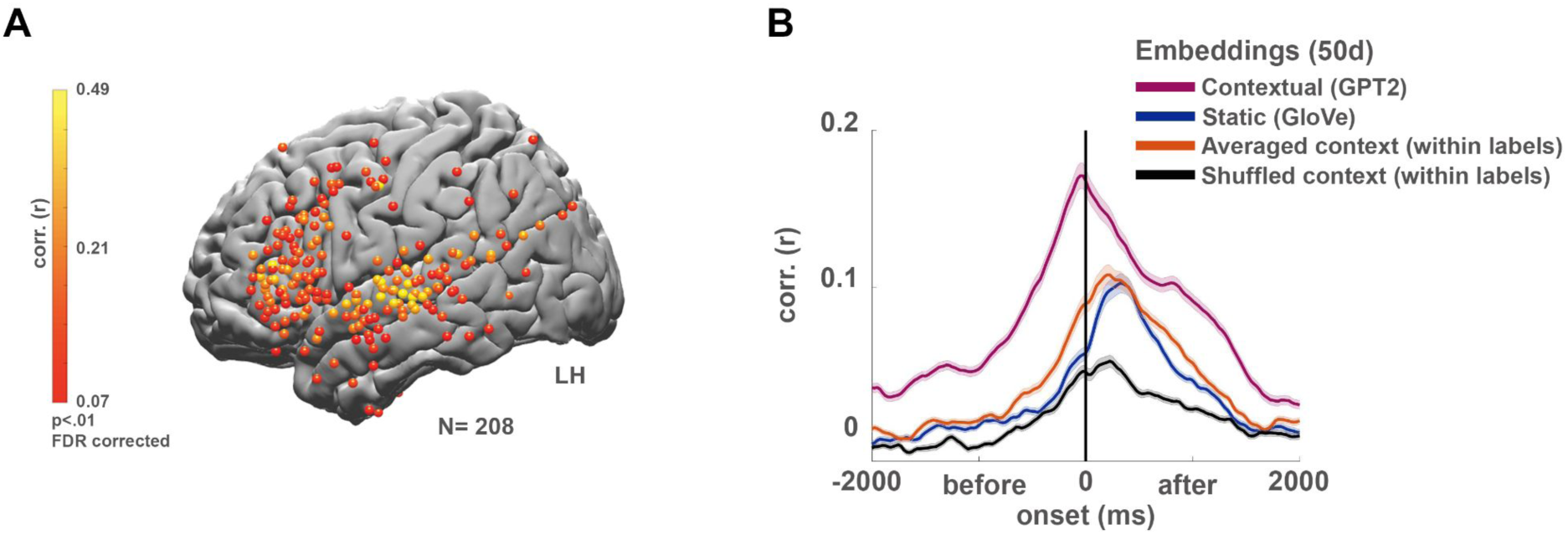
Contextual (GPT-2) embeddings improve the modeling of neural responses before word onset. **A)** Peak correlation between predicted and actual word responses for the contextual (GPT-2) embeddings. Using contextual embeddings significantly improved the encoding model’s ability to predict the neural signals for unseen words across many electrodes. **B)** Encoding model performance for contextual embeddings (GPT-2) aggregated across 160 electrodes with significant encoding for GloVe (Fig. 3C): contextual embeddings (purple), static embeddings (GloVe, blue), contextual embeddings averaged across all occurrences of a given word (orange), contextual embeddings shuffled across context-specific occurrence of a given word (black).

#### Modeling the context versus predicting the upcoming word

The improved ability to predict neural responses before word onset using contextual embedding can be attributed to two related factors that are absent in the static word embeddings (e.g., GloVe): 1) the brain, like GPT-2, aggregate information about the preceding words in the story into contextual embeddings; and 2) GPT-2 embeddings contain additional predictive information, not encoded in static embeddings, about the identity of the upcoming word in the sequence. By carefully manipulating the contextual embeddings and developing an embedding-based decoder, we show how both context and next-word prediction contribute to contextual embeddings’ improved ability to model the neural responses.

#### Representing word meaning in unique contexts

Going above and beyond the information encoded in GloVe, GPT-2’s capacity for representing context captures additional information in neural responses. A simple way to represent the context of prior words is to combine (i.e., concatenating) the static embeddings of the preceding sequence of words. To test this simpler representation of context, we concatenated GloVe embeddings for the ten preceding words in the text into a longer “context” vector and compared the encoding model performance to GPT-2’s contextual embeddings (after reducing both vectors to 50 dimensions using PCA). While the concatenated static embeddings were better in predicting the prior neural responses than the original GloVe vectors (which only capture the current word), they still underperformed GPT-2’s encoding before word articulation (Fig. S9). This result suggests that GPT-2’s contextual embeddings are better suited to compress the contextual information embedded in the neural responses than static embeddings (even when concatenated).

A complementary way to demonstrate that contextual embeddings uncover aspects of the neural activity that static embeddings cannot capture is to remove the unique contextual information from GPT-2 embeddings. We removed contextual information from GPT-2’s contextual embeddings by averaging all embeddings for each unique word (e.g., all occurrences of the word “monkey”) into a single vector. This analysis was limited to words that have at least 5 repetitions (see Material and Method).Thus, we collapsed the contextual embedding into a static embedding in which each unique word in the story is represented by one unique vector. The resulting embeddings are still specific to the overall topic of this particular podcast (unlike GloVe). Still, they do not contain the local context for each occurrence of a given word (e.g., the context in which “*monkey*” was used in sentence 5 versus the context in which it was used in sentence 50 of the podcast). Indeed, removing context from the contextual embedding by averaging the vector for each unique word effectively reduced the encoding model’s performance to that of the static GloVe embeddings (Fig. 6B, orange).

Finally, we examined how the specificity of the contextual information in the contextual embeddings improved the ability to model the neural responses to each word. To that end, we scrambled the embeddings across different occurrences of the same word in the story (e.g., switched the embedding of the word “*monkey*” in sentence 5 with the embedding for the word “*monkey*” in sentence 50). This manipulation tests whether contextual embeddings are necessary for modeling neural activity for a specific sequence of words. Scrambling the same word occurrences across contexts substantially reduced the encoding model performance (Fig. 6B, black), pointing to the contextual dependency represented in the neural signals. Taken together, these results suggest that contextual embeddings provide us with a new way to model the context-dependent neural representations of words in natural contexts.

#### Using contextual embeddings for predicting the next word from neural responses

Finally, we apply a **decoding** analysis to demonstrate that in addition to better modeling the neural responses to context, contextual embeddings also improve our ability to read information from the neural responses as to the identity of upcoming words. This demonstrates the duality of representing the context and the next-word prediction in the brain.

The **encoding** model finds a mapping from the embedding space to the neural responses that is used during the test phase for predicting neural responses. The **decoding** analysis inverts this procedure to find a mapping from neural responses, across multiple electrodes and time points, to the embedding space.^52^. This decoding analysis provides complementary insights to the encoding analysis by aggregating across electrodes and quantifies how much predictive information about each word’s identity is embedded in the spatiotemporal neural activity patterns before and after word-onset.

The decoding analysis was performed in two steps. First, we trained a deep convolutional neural network to aggregate neural responses (Fig. 7A and Appendix I) and map this neural signal to the arbitrary, static (GloVe based) and to the contextual (GPT-2 based) embedding spaces (Fig. 7B). In order to conservatively compare the performance of GPT-2 based embedding to GloVe embedding, we used as input the signal from the electrodes that were found significant for GloVe based encoding. To further ensure that the decoding results are not affected by the electrodes selection procedure, for each test fold, we selected the electrodes using the remaining 80% of the data (see Materials and Methods). In order to get a reliable estimation of accuracy per word-label we included words with at least 5 repetitions, which included 69% of the words in the story (for the full list of words see Appendix II). Second, the predicted word embeddings were used for word classification based on their cosine distance from all embeddings in the dataset (Fig. 7C). Although we evaluated the decoding model using classification, the classifier predictions were constrained to rely only on the embedding space’s information. This is a more conservative approach than an end-to-end word classification, which may capitalize on acoustic information in the neural signals that are not encoded in the language models.

**Figure 7.**
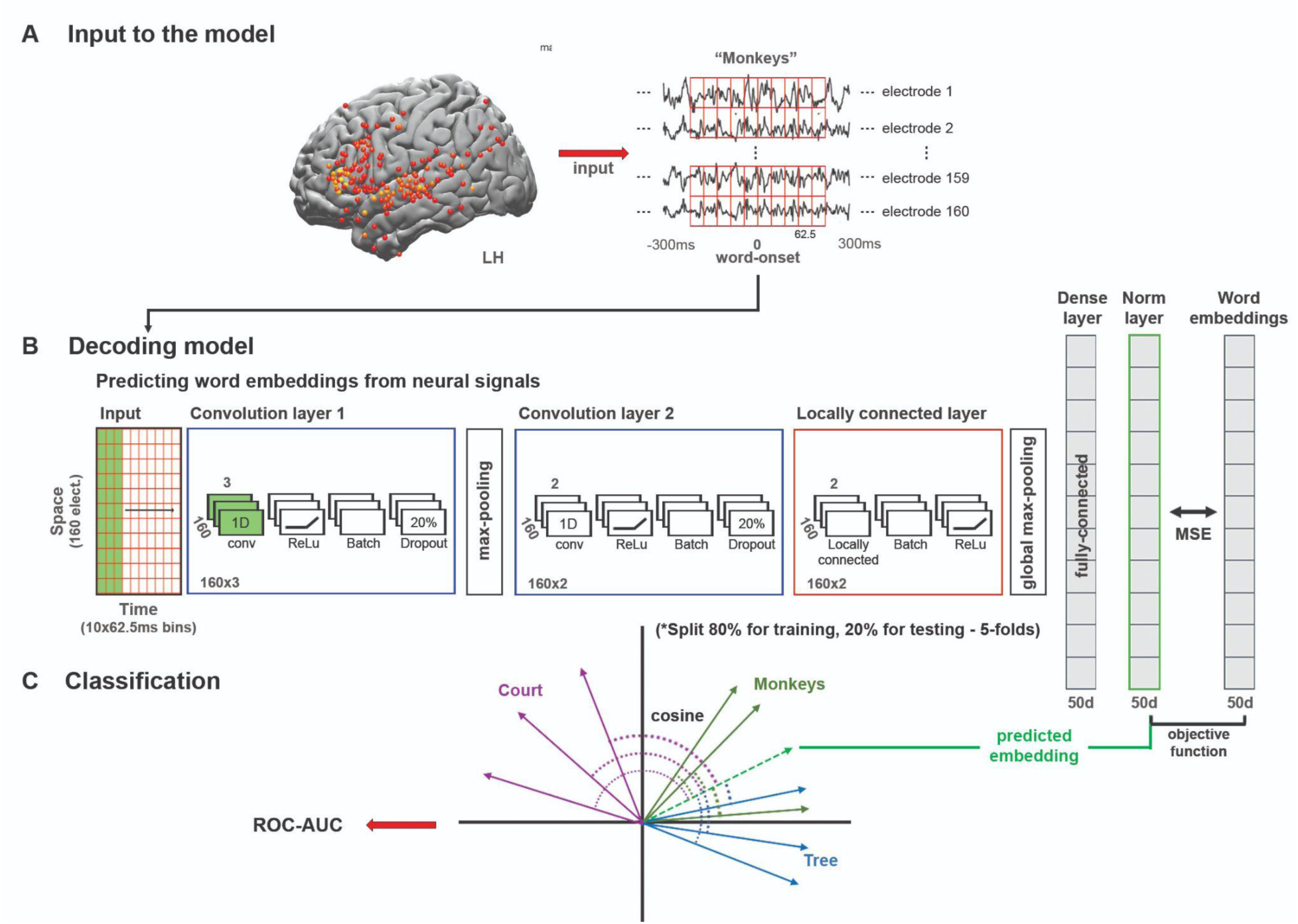
Deep nonlinear decoding model used to predict words from neural responses before and after word-onset. **A)** Neural data from left hemisphere electrodes with significant encoding model performance using GloVe embeddings were used as input to the decoding model. For each fold, electrodes selection was performed on 80% of the data that were not used for testing the model. The stimulus is segmented into individual words and aligned to the brain signal at each lag. **B)** Schematic of the feedforward deep neural network model that learns to project the neural signals for the words into the arbitrary embedding, static semantic embedding (GloVe) or contextual embedding (GPT-2) space (for full description, see Appendix I). The input (currently represented as 160X10 matrix), changes its dimensions for each of the five folds based on the number of significant electrodes for each fold. The model was trained to minimize the mean squared error (MSE) when mapping the neural signals into the embedding space. **C)** The decoding model was evaluated using a word classification task. The quality of word classification is based on the embedding space used to construct ROC-AUC scores. This enables us to assess how much information about specific words is extractible from the neural activity via the linguistic embedding space.

Using a contextual decoder greatly improved our ability to classify words’ identity over decoders relying on static or arbitrary embeddings (Fig. 8). We evaluated classification performance using the area under the receiver operating characteristic curve (ROC-AUC). A model that only learns to use word frequency statistics (e.g., only guessing the most frequent word) will result in a ROC-AUC curve that falls on the diagonal line (AUC = 0.5) suggesting that the classifier does not discriminate between the words^53^. Classification using GPT-2 (average AUC of 0.74 for lag 150) outperformed GloVe and arbitrary embeddings (average AUC of 0.68 and 0.68, respectively) before and after word-onset. To compare the performance of the classifiers based on GPT-2 and GloVe at each lag, we performed a paired-sample t-test between the AUCs of the words in the two models. Each unique word (class) in each lag has an AUC value computed from the GloVe based model, and an AUC value computed from the GPT-2 based model. The results were corrected for multiple tests by controlling the false discovery rate (FDR)^54^.

**Figure 8.**
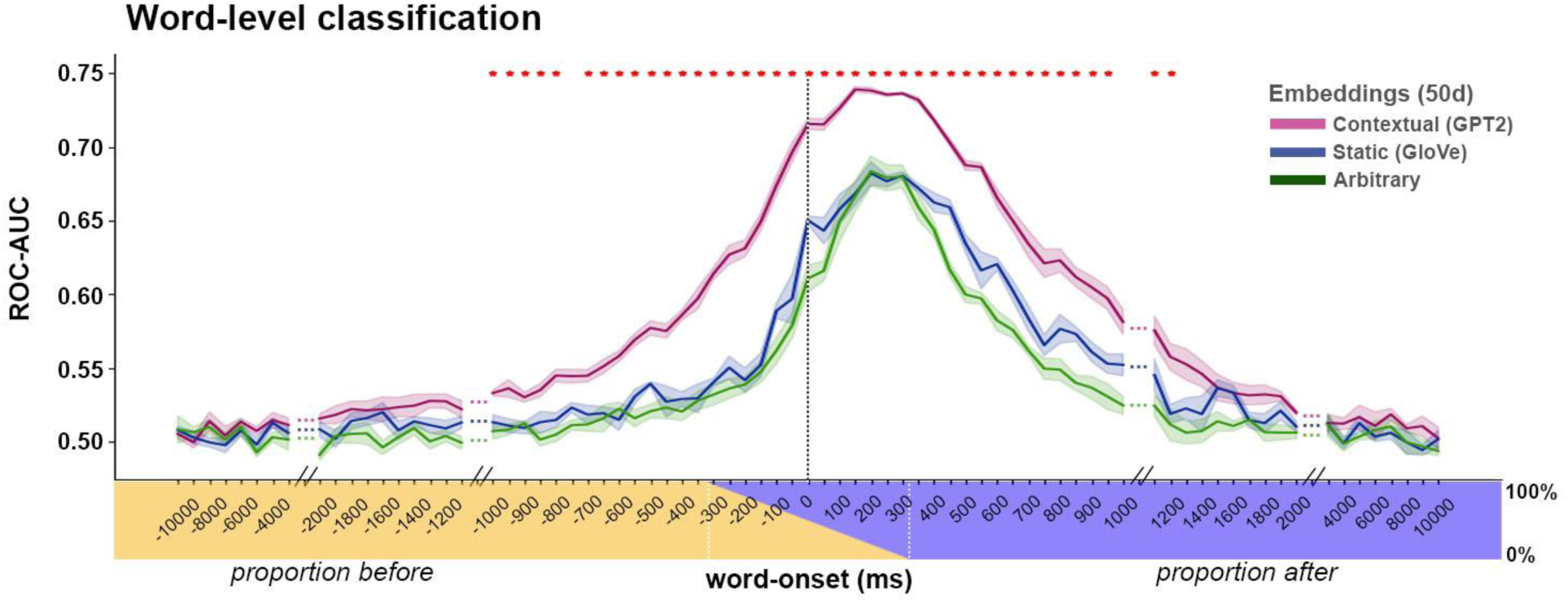
Using a decoding model for classification of words before and after word-onset. *Word-level classification.* Classification performance for contextual embeddings (GPT2; purple), static embeddings (GloVe; blue), and arbitrary embeddings (green). The averaged values are weighted by the frequency of the words in the test set. The *x*-axis labels indicate the center of each 625 ms window used for decoding at each lag (between -10 to 10 sec). The colored stripe indicates the proportion of pre-(yellow) and post- (blue) word onset time points in each lag. Error bars denote SE across five test folds. Note that contextual embeddings improve classification performance over GloVe both before and after word-onset. Significance was assessed using paired-sample t-test of the AUCs for each unique word, comparing the AUCs of the GloVe-based decoding and GPT2-based decoding. The comparison was done for each leg separately and results were FDR corrected (q < .01).

A closer inspection of the GPT-2-based decoder indicates that the classifier managed to detect reliable information about the identity of words several hundred milliseconds before word onset (Fig. 8). In particular, starting at about -1000 ms before word onset, when the neural signals were integrated across a window of 625 ms, the classifier detected predictive information about the next word’s identity. The information about the next word’s identity gradually increased and peaked at an average AUC of 0.74 at a lag of 150 ms after word onset, when the signal was integrated across a window from -162.5 ms to 462.5 ms. GloVe embeddings show a similar trend with a marked reduction in classifier performance (Fig. 8, blue). The decoding model’s capacity to classify words before word onset demonstrates that the neural signal contains a considerable amount of predictive information about the meaning of the next word, up to a second before it is perceived. At long time scales (more than 2 seconds), all decoders’ performance dropped to chance.

## Discussion

Deep language models (DLMs) provide a new modeling framework that drastically departs from classical psycholinguistic models. They are not designed to learn a concise set of interpretable syntactic rules to be implemented in novel situations, nor do they rely on part of speech concepts or other linguistic terms. Rather, they learn from surface-level linguistic behavior to predict and generate the contextually appropriate linguistic outputs. The current paper provides compelling behavioral and neural evidence for shared computational principles between the way the human brain and autoregressive DLMs process natural language.

### Spontaneous prediction as a keystone of language processing

Autoregressive DLMs learn according to the simple self-supervised objective of context-based next-word prediction. The extent to which humans are spontaneously engaged in next-word predictions as they listen to continuous, minutes-long, natural speech has been underspecified. Our behavioral results revealed a robust capacity for next-word prediction in real-world stimuli, which matches a modern autoregressive DLM (Fig. 2). Neurally, our findings demonstrate that the brain constantly and actively represents forthcoming words in context during listening to natural speech. The predictive neural signals are robust, and can be detected hundreds of milliseconds before word-onset. Notably, the next-word prediction processes are associated with listeners’ contextual expectations and can be dissociated from the processing of the actually perceived words after word-onset (Fig. 4B, S8).

### Pre-onset predictions are coupled with post-onset surprise signals

Autoregressive DLMs (as GPT-2) provide a unified computational framework that connects pre-onset word prediction with post-onset surprise (error signals). Our behavioral results indicate that human and GPT-2 predictions match during the processing of natural language in real-world contexts. Furthermore, the results show that we can rely on GPT-2’s internal pre-onset confidence (uncertainty) in its predictions (entropy) and post-onset surprise (cross-entropy) to model the brain’s internal neural activity as it processes language. The correlations between the surprise induced by the DLM and the neural signals corroborate the observation that DLMs can be used to predict surprise related ERPs^55^. The correlational nature of the result, however, calls for future research that better establishes a causal link between the entropy and surprise of the models and the neural response ^56–58^.

The enhanced encoding (Fig. 4B) and enhanced neural activity (Fig. 5A) for surprising words 400 ms after word onset can be associated with different cognitive processes: estimating the surprise level, the allocation of additional attentional resources to process the surprising word, as well as additional processes needed to extract the semantic meaning of unpredictable words. Future research will be needed to fully characterize the post-onset surprise signals.

### Context-specific meaning as a keystone of language processing

As each word attains its full meaning in the context of preceding words over multiple timescales, language is fundamentally contextual ^42, 59^. Even a single change to one word or one sentence at the beginning of a story can alter the neural responses to all subsequent sentences ^48, 60^. We demonstrated that the contextual word embeddings learned by DLMs provide a new way to compress linguistic context into a numeric vector space, which outperforms the use of static semantic embeddings (Figs. 6B, 8, S4, S9). While static embeddings and contextual embeddings are fundamentally different, our neural results also hint at how they relate to each other. Our results indicate that both static and contextual embeddings can predict neural responses to single words in many language areas^16^ along the superior temporal cortex, parietal lobe, and inferior frontal gyrus. Switching from static to contextual embeddings boosted our ability to model neural responses during natural speech processing across many of these brain areas. Finally, averaging contextual embeddings associated with a given word, removed the contextual information, and changed GPT-2’s contextual embedding back into static word embeddings (Fig. 6B). Taken together, these results suggest that the brain is coding for the semantic relationship among words contained in static embeddings while also being tuned to the unique contextual relationship between the specific word and the preceding words in the sequence^61^.

### Using autoregressive language model as a cognitive model

In this paper, we describe three shared computational principles that reveal a strong link between the way the brain and DLMs process natural language. These shared computational principles, however, do not imply that the human brain and DLMs implement these computations in a similar way ^62,63,64,65^. For example, many state-of-the-art DLMs rely on transformers, a type of neural network architecture developed to solve sequence transduction. Transformers are designed to parallelize a task that is largely computed serially, word by word, in the human brain. Therefore, while transformer models are an impressive engineering achievement, they are not biologically feasible. Many other ways, however, are possible to transduce a sequence into a contextual embedding vector. To the extent that the brain relies on a next-word prediction objective to learn how to use language in context, it likely uses a different implementation^64^.

### Psycholinguistic models versus deep language models

DLMs were engineered to solve a fundamentally different problem than psycholinguistic language models. Psycholinguistic language models aim to uncover a set of generative (learned or innate) rules to be used in infinite, novel situations ^66^. Finding a set of linguistic rules, however, was found to be challenging. There are numerous exceptions for every rule, conditioned by discourse context, meaning, dialect, genre, and many other factors ^59, 67–71^. In contrast, DLMs aim to provide the appropriate linguistic output given the prior statistics of language use in similar contexts ^20, 72^. In other words, psycholinguistic theories aim to describe observed language in terms of a succinct set of explanatory constructs. DLMs, in contrast, are performance-oriented and are focused on learning how to generate formed linguistic outputs as a function of context, while deemphasizing interpretability ^73^. The reliance on performance creates an interesting connection between DLMs and usage- (context-) based constructionist approaches to language ^67, 70, 74^. Furthermore, DLMs avoid the circularity built into many psycholinguistic language models that rely on linguistic terms to explain how language is encoded in neural substrates ^19, 75^. Nevertheless, the internal contextual embedding space in DLMs can capture many aspects of the latent structure of human language, including syntactic trees, voice, co-references, morphology, and long-range semantic and pragmatic dependencies ^1,76–78^. This discussion demonstrates the power (over the more traditional approaches) of applying brute-force memorization and interpolation for learning how to generate the appropriate linguistic outputs in light of prior contexts^20^.

Observational work in developmental psychology suggests that children are exposed to tens of thousands of words in contextualized speech each day, creating a large data volume available for learning ^79–81^. The capacity of DLMs to learn language relies on gradually exposing the model to millions of real-life examples. Our finding of spontaneous predictive neural signals as participants listen to natural speech suggests that active prediction may underlie humans’ lifelong language learning. Future studies, however, will have to assess whether these cognitively plausible, prediction-based feedback signals are at the basis of human language learning and whether the brain is using such predictive signals to guide language acquisition. Furthermore, as opposed to autoregressive DLMs, it is likely that the brain relies on additional simple objectives at different timescales to facilitate learning ^20, 82^.

## Conclusion

This paper provides evidence for three shared core computational principles between deep language models and the human brain. While DLMs may provide a building block for our high-level cognitive faculties, they undeniably lack certain central hallmarks of human cognition. Linguists were primarily interested in how we construct well-formed sentences, exemplified by the famous grammatically correct but meaningless sentence composed by Noam Chomsky “colorless green ideas sleep furiously”^2^. Similarly, DLMs are generative in the narrow linguistic sense of being able to generate new sentences that are grammatically, semantically, and even pragmatically well-formed at a superficial level. However, although language may play a central organizing role in our cognition, linguistic competence is insufficient to capture thinking. Unlike humans, DLMs cannot think, understand, or generate new meaningful ideas by integrating prior knowledge. They simply echo the statistics of their input^83^. Going beyond the importance of language as having a central organizing role in our cognition, DLMs indicate that linguistic competence may be insufficient to capture thinking. A core question for future studies in cognitive neuroscience and machine learning is how the brain can leverage predictive, contextualized linguistic representations, like those learned by DLMs, as a substrate for generating and articulating new thoughts.

## Acknowledgements

We thank Adele Goldberg, Rita Goldstein, Sebastian Michelmann, Meir Meshulam, Manoj Kumar, Malcolm Slaney, and Alex Huth for technical and conceptual assistance that motivated and informed this manuscript’s writing. This work was supported by the National Institutes of Health under award numbers DP1HD091948 (A.G, Z.Z, A.P, B.A, G.C, A.R, C.K, F.L, A.F and U.H.), R01MH112566 (S.A.N.), NIH R01NS109367-01 to A.F, Finding A Cure for Epilepsy and Seizures (FACES), and DataX Fund, Schmidt Futures Foundation.

## Materials and Methods

### Transcription and alignment

Stimuli for the behavioral test and ECoG experiment were extracted from a 30-minute story “So a Monkey and a Horse Walk Into a Bar: Act One, Monkey in the Middle” taken from the ”This American Life podcast”. The story was manually transcribed and aligned to the audio by marking the onset and offset of each word. Sounds such as laughter, breathing, lip-smacking, applause, and silent periods were also marked to improve the alignment’s accuracy. The audio was downsampled to 11 kHz and the Penn Phonetics Lab Forced Aligner was used to automatically align the audio to the transcript^84^. The forced aligner uses a phonetic Hidden Markov model to find the temporal onset and offset of each word and phoneme in the story. After automatic alignment was complete, the alignment was manually evaluated by an independent listener..

### Behavioral word-prediction experiment

To obtain a continuous measure of prediction, we developed a novel sliding-window behavioral paradigm where healthy adult participants made predictions for each upcoming word in the story. 300 participants completed a behavioral experiment on Mechanical Turk. Workers were native English speakers with a proven record of successfully completing at least 95% of a minimum of 500 prior tasks on the platform. Since predicting each word in a 30-minute (5113 words) story is taxing, we divided the story into six segments and recruited six non-overlapping groups of 50 participants to predict every upcoming word within each segment of the story (about 830 words per group of participants). The first group was exposed to the first two words in the story and then asked to predict the upcoming (i.e., third) word. After entering their prediction, the actual next word was revealed, and participants were asked again to predict the next upcoming (i.e., fourth) word in the story. Once 10 words were displayed on the screen, the left-most word was removed and the next word was presented (Fig. 2B). The procedure was repeated, using a sliding window until the first group provided predictions for each word in the story’s first segment. Each of the other five groups listened uninterruptedly to the prior segments of the narrative and started to predict the next word at the beginning of their assigned segments.

Next, we calculated a mean prediction performance (proportion of participants predicting the correct word) across all 50 listeners for each word in the narrative, which we refer to as the “predictability score” (Fig. 2C). A predictability score of 1 indicates that all subjects correctly guessed the next word, and a predictability score of 0 indicates that no participant predicted the upcoming word. Due to a technical error, data for 33 words were omitted, and thus the final data contained 5078 words. Importantly, before calculating the scores we used Excel’s spell checker to locate and correct spelling mistakes.

### N-gram models

We trained 2- to 5-gram models using the NLTK python package and its built-in “Brown” corpus ( http://www.nltk.org/nltk_data/ ). All punctuations were removed and letters lower-cased. We trained separate models using no-smoothing, Laplace smoothing or Kneser-Ney smoothing. Then we used each model to extract the probability of a word given its preceding n – 1 context in the podcast transcript. We also extracted the most likely next word prediction to compare agreement with human responses. In terms of correlations to human behavior, no-smoothing probabilities were comparable to ones achieved by applying different smoothing methods.

### ECoG experiment

Ten patients (5 female; 20–48 years old) listened to the same story stimulus from beginning to end. Participants were not explicitly made aware that we would be examining word prediction in our subsequent analyses. One patient was removed from further analyses, due to excessive epileptic activity and low SNR across all experimental data collected during the day. All patients experienced pharmacologically refractory complex partial seizures and volunteered for this study via the New York University School of Medicine Comprehensive Epilepsy Center. All participants had elected to undergo intracranial monitoring for clinical purposes and provided oral and written informed consent before study participation, according to the New York University Langone Medical Center Institutional Review Board. Patients were informed that participation in the study was unrelated to their clinical care and that they could withdraw from the study at any point without affecting their medical treatment.

For each patient, electrode placement was determined by clinicians based on clinical criteria (Fig. 3A). One patient consented to have an FDA-approved hybrid clinical-research grid implanted which includes standard clinical electrodes as well as additional electrodes in between clinical contacts. The hybrid grid provides a higher spatial coverage without changing clinical acquisition or grid placement. Across all patients, a total of 1106 electrodes were placed on the left hemisphere and 233 on the right hemisphere. Brain activity was recorded from a total of 1339 intracranially implanted subdural platinum-iridium electrodes embedded in silastic sheets (2.3 mm diameter contacts, Ad-Tech Medical Instrument; for the hybrid grids 64 standard contacts had a diameter of 2 mm and additional 64 contacts were 1 mm diameter, PMT corporation, Chanassen, MN). Decisions related to electrode placement and invasive monitoring duration were determined solely on clinical grounds without reference to this or any other research study. Electrodes were arranged as grid arrays (8 × 8 contacts, 10 or 5 mm center-to-center spacing), or linear strips. Altogether, the subdural electrodes covered extensive portions of lateral frontal, parietal, occipital, and temporal cortex of the left and/or right hemisphere (Fig. 3A for electrode coverage across all subjects).

Recordings from grid, strip and depth electrode arrays were acquired using one of two amplifier types: NicoletOne C64 clinical amplifier (Natus Neurologics, Middleton, WI), bandpass filtered from 0.16–250 Hz, and digitized at 512 Hz; Neuroworks Quantum Amplifier (Natus Biomedical, Appleton, WI) recorded at 2048 Hz, highpass filtered at 0.01 Hz and then resampled to 512 Hz.

Intracranial EEG signals were referenced to a two-contact subdural strip facing towards the skull near the craniotomy site. All electrodes were visually inspected, and those with excessive noise artifacts, epileptiform activity, excessive noise, or no signal were removed from subsequent analysis (164/1065 electrodes removed).

Pre-surgical and post-surgical T1-weighted MRIs were acquired for each patient, and the location of the electrodes relative to the cortical surface was determined from co-registered MRIs or CTs following the procedure described by Yang and colleagues^85^. Co-registered, skull-stripped T1 images were nonlinearly registered to an MNI152 template and electrode locations were then extracted in Montreal Neurological Institute (MNI) space (projected to the surface) using the co-registered image. All electrode maps are displayed on a surface plot of the template, using the Electrode Localization Toolbox for MATLAB available at (https://github.com/HughWXY/ntools_elec).

### Preprocessing

Data analysis was performed using the FieldTrip toolbox^86^, along with custom preprocessing scripts written in MATLAB 2019a (MathWorks). 66 electrodes from all patients were removed due to faulty recordings. The analyses described are at the electrode level. Large spikes exceeding 4 quartiles above and below the median were removed and replacement samples were imputed using cubic interpolation. We then re-referenced the data to account for shared signals across all channels using either the Common Average Referencing (CAR) method ^86, 87^ or an ICA-based method^88^ (based on the participant’s noise profile). High-frequency broadband (HFBB) power frequency provided evidence for a high positive correlation between local neural firing rates and high gamma activity^89^. The high gamma band fluctuation exhibited good estimations in the neural spiking population near each electrode^90^.

Broadband power was estimated using 6-cycle wavelets to compute the power of the 70-200 Hz band, excluding 60, 120, 180 Hz line noise. Power was further smoothed with a Hamming window with a kernel size of 50 ms. To preserve the temporal structure of the signal, we used zero-phase symmetric filters. The estimate of the broadband power using wavelets and symmetric filters, by construction, induces some temporal uncertainty, given that information over tens of milliseconds is combined. The amount of temporal uncertainty, however, is small relative to the differences between pre-onset and post-onset effects reported in the paper. First, as the wavelet computation was done using six cycles and the lower bound of the gamma-band was 70 Hz, the wavelet computation introduces a 6/70-s uncertainty window centered at each time point. Thus, there is a leak from no more than 43 ms of future signal to data points in the preprocessed signal. Second, the smoothing procedure applied to the broadband power introduces a leak of up to 50 ms from the future. Overall the leak from the future is at max 93 ms. As recommended by^91^ this was empirically verified by examining the preprocessing procedure on an impulse response (showing a leak of up to ∼90 ms, see Fig. S10).

A scheme of the procedure:

#### Despike

● Removing recordings that deviate more than 3 times the interquartile from the mean value of the electrode..
● Interpolating the removed values using cubic interpolation.

#### Detrend

● Common average referencing / Remove ICA components.

#### Broadband power

● Using 6-cycle wavelets to compute the power of the 70-200 Hz band, excluding 60,180 Hz.
● Natural log transformation
● Z-score transformation

#### Temporal smoothing

● Using a filter to smooth the data with a Hamming window with kernel size of 50 ms. The filter was applied in both the forward and reverse directions in order to maintain the temporal structure, specifically the encoding peak-onset (zero-phase).

### Encoding analysis

In this analysis, a linear model is implemented for each lag for each electrode relative to word-onset, and is used to predict the neural signal from word embeddings (Fig. 3B). The calculated values are the correlations between the predicted signal and the held out actual signal at each lag (separately for each electrode), indicating the linear model’s performance. Before fitting the linear models for each time point, we implement a running window averaging across a 200 ms window. We assess the linear models’ performance (model for each lag) in predicting neural responses for held-out data using a 10-fold cross-validation procedure. The neural data were randomly split into a training set (i.e., 90% of the words) for model training and a testing set (i.e., 10% of the words) for model validation. On each fold of this cross-validation procedure, we used ordinary least-squares multiple linear regression to estimate the regression weights from 90% of the words. We then applied those weights to predict the neural responses to the other 10% of the words. The predicted responses for all ten folds were concatenated so a correlation between the predicted signal and actual signal was computed over all the words of the story. This entire procedure was repeated at 161 lags from -2000 ms to 2000 ms in 25 ms increments relative to word onset.

Part of the encoding analysis involves selection of words to include in the analysis. For each analysis we included the relevant words. Fig. 4A includes all the words in the transcription that have a GloVe embedding (excluding non-words like “ah”, “um”, etc.), totaling 4843 words (1190 unique words). Fig. 4B-D comprises 2886 accurately predicted words (796 unique words) and 1762 inaccurately predicted words (562 unique words). Lastly, Fig. 6B comprises 3279 words (165 unique words) that have both GloVe and GPT-2 embeddings, to allow for comparison between the two, and at least five repetitions for the average context and shuffle context conditions.

### Accuracy split

To model the brain’s prediction, the podcast’s transcription words were split into two groups. Each word was marked whether it was one of the top-5 most probable words in the distribution that GPT-2 predicted given its past context (up to 1024 previous tokens) or not (see Fig. 1). 62% of the words included in the top-5 predicted words given their context and were classified as correctly predicted using this accuracy measure. The other words were classified as incorrectly predicted (38%). To control possible confounds that stem from the accurately predicted words that are bigger than the inaccurately predicted group, we also report results from classifying the words according to top-1 probability in Fig. S8. Using the top-1 measure for human prediction, we got a group of 36% correctly predicted words.

### Confidence and surprise measures

We associate pre-onset neural activity with confidence in prediction, and post-onset neural activity with surprise (prediction-error). Both could be estimated using the GPT-2 language model. Given a sequence of words, autoregressive DLMs (such as GPT-2) induce a distribution of probabilities for the next possible word (i.e., predict the next word). We use the entropy of this distribution as a measure for the confidence in prediction ^14, 22, 46^ :

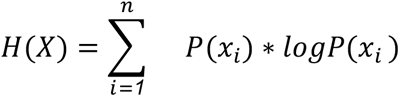

Where *n* is the vocabulary size and P(xi) is the probability (assigned by the model) of the i-th word in the vocabulary. It is intuitively understood as the distance from a uniform distribution ^92^. The closer a distribution is to the uniform distribution, the less confident its predictions are, the **higher** the entropy is. In other words, the more confident the model is, the more concentrated the predictions are on a small number of words (with higher probabilities). Thus, the more confident the model is, the **lower** the entropy is.

To estimate the surprise we use the cross-entropy measure. Cross-entropy is the loss function used to attenuate the autoregressive DLMs weights, given its predictions (i.e., the distribution) and the actual word. The lower the probability of the actual word before its onset, the higher the surprise it induces. It is defined by:

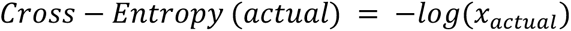

To be concise, entropy and cross-entropy (or surprise) are closely related. While entropy represents the distance of a distribution from the uniform distribution, the cross-entropy describes the distance between the distribution to the 1-hot distribution. In the 1-hot distribution, all the vocabulary words are assigned with a probability of 0, except the actual next word in the sequence. Thus, canceling all the words but one in the entropy formula yields the cross-entropy definition.

### Significance tests

To identify significant electrodes, we used a randomization procedure. At each iteration, we randomized each electrode’s signal phase uniform distribution, thus disconnecting the relationship between the words and the brain signal but preserving the autocorrelation in the signal^93^. Then we performed the entire encoding procedure for each electrode. We repeated this process 5000 times. After each iteration, the encoding model’s maximal value across all 161 lags was retained for each electrode. We then took the maximum value for each permutation across electrodes. This resulted in a distribution of 5000 values, which was used to determine significance for all electrodes. For each electrode a *p*-value (Fig. 3C, 6A, S3 & S4) was computed as the percentile of the non-permuted encoding model’s maximum value across all lags from the null distribution of 5000 maximum values. Performing a significance test using this randomization procedure evaluates the null hypothesis that there is no systematic relationship between the brain signal and the corresponding word embedding. This procedure yielded a *p*-value per electrode. To correct for multiple electrodes we used false-discovery-rate (FDR^54^). Electrodes with *q*-values less than .01 were considered significant.

To test each lag’s significance for two different encoding models for the same group of electrodes (Figs. 4ABD, S5, S8 & S9) we used a permutation test. Each electrode has encoding values for two encoding models. We randomly swapped the assignment of the encoding values between the two models (50% of the pairs were swapped). Then we computed the average of the pairwise differences to generate a null distribution at each lag.To account for multiple tests across lags, we adjusted the resulting *p*-values to control the false discovery rate^54^. A threshold was chosen to control the FDR at *q*=.01.

To set a threshold above which average encoding values are significant (Figs. 4A, S6 & S7), we used a bootstrapping^94^. For each bootstrap, a sample matching the subset size was drawn with replacement from the encoding performance values for the subset of electrodes. The mean of each bootstrap sample was computed. This resulted in a bootstrap distribution of 5000 mean performance values for each lag. The bootstrap distribution was then shifted by the observed value to perform a null hypothesis test^94^. To account for multiple tests across lags, we adjusted the resulting *p*-values to control the false discovery rate^54^. A threshold was chosen to control the FDR at *q*=.01.

For identifying electrodes that demonstrate pre-onset prediction, post-onset prediction error (Fig. 4C), we conducted the following: For pre-onset prediction we did a similar permutation test as described above for Fig. 3C; however, we looked only at lags between -500 ms to -100 ms before onset. Because the window size of the encoding is 200 ms, limiting to an upper bound of -100 ensures we are not looking at data post-onset. For post-onset prediction error we did a similar bootstrapping as described for Fig. 4BD, however, we limited the range of the significance testing to 350 ms to 450 ms post onset (around N400).

To statistically assess the pre-onset prediction for neural responses to correctly predicted words (Fig. 5), we did a permutation test (such as the one described for 4ABD), however, we were also constrained to lags at which the neural responses were significant on their own (not with respect to the neural response of the inaccurate conditional brain response). The same procedure was implemented for the significant test of post-onset surprise.

### Contextual embedding extraction

We extracted contextualized word embeddings from GPT-2 for our analysis. We used the pre-trained version of the model implemented in the Hugging Face environment ^95^. We first converted the words from the raw transcript (including punctuation and capitalization) to tokens which were either whole words or sub-words (there’s -> there ’s). We used a sliding window of 1024 tokens, moving one token at a time, to extract the embedding for the final word in the sequence (i.e. the word and its history). Encoding these tokens into integer labels, we then fed them into the model, and in return, we received the activations at each layer in the network (also known as a hidden state). GPT-2 has 48 layers, but we focused only on the final one, before the classification layer. Finally, the token of interest was the final word of the sequence, yet we used the second-to-last token as the hidden state for the last word because it was the same activation embedding that was used to predict that word. With embeddings for each word in the raw transcript, we aligned this list with our spoken-word transcript that did not include punctuation, thus retaining only full words.

### Decoding analysis

The goal of this analysis was to predict words from the neural signal. The input neural data were averaged in 10 62.5-ms bins spanning 625 ms for each lag. Each bin consisted of 32 data points (the neural recording sampling rate was 512Hz).

The neural network decoder (see architecture in Appendix I) was trained to predict a word’s embedding from the neural signal at a specific lag. The data was split into 5 non-overlapping temporal intervals (i.e., folds) and used in a cross-validation procedure. Each fold consisted of a mean of 717.04 training words (SD = 1.32). Three folds were used for training the decoder (training set), one fold was used for early stopping (development set), and one fold was used to assess model generalization (test set). The neural net was optimized to minimize the MSE when predicting the embedding. The decoding performance was evaluated using a classification task assessing how well the decoder can predict the word label from the neural signal. We used the receiver operating characteristic curve (ROC-AUC) measure.

To ensure that the decoding ability is not affected by the electrode selection procedure we used the training and validation folds (80% of the data) to choose the electrodes for each model. We used the same significance test used to locate GloVe-based significant encodings (Fig. 3C). This procedure yielded a different number of electrodes for each model, ranging between 114 to 132.

To calculate the ROC-AUC, we computed the cosine distance between each of the predicted embeddings and the embeddings of all instances of each unique word label. The distances were averaged across unique word labels, yielding one score for each word label (i.e., logit). We used a softmax transformation on these scores (logits). For each label (classifier), we used the logits and the information of whether the instance matched the label to compute a ROC-AUC for the label. We plotted the weighted ROC-AUC according to the word’s frequency in the test set . In order to get reliable ROC-AUC scores we chose words with at least 5 repetitions in the training set (69% of the overall words in the narrative; see Appendix II for word list).

To improve the performance of the decoder, we implemented an ensemble of models. We independently trained 10 decoders with randomized weight initializations and randomized the batch order fed into the neural net for each lag. This procedure generated 10 predicted embeddings. Thus, for each predicted embedding, we repeated the distance calculation from each word label 10 times. These 10 values were averaged and later used for ROC-AUC.

## Supplementary Information

**Figure S1.**
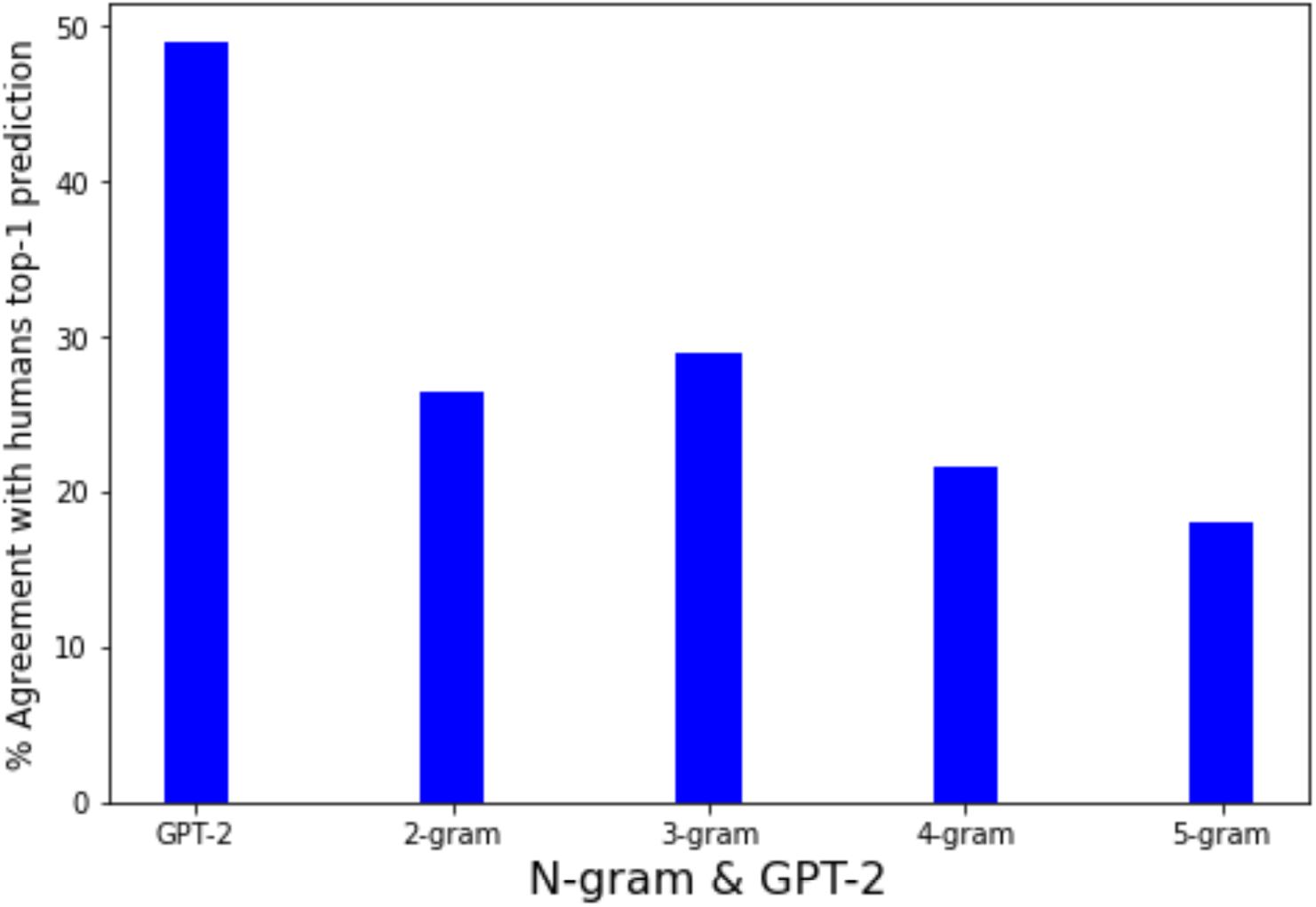
Comparing agreement with human prediction between most probable predictions based on n-grams or GPT-2. The plots show higher agreement between human predictions and GPT-2‘s top-1 predictions than all the n-gram model predictions we trained.

**Figure S2.**
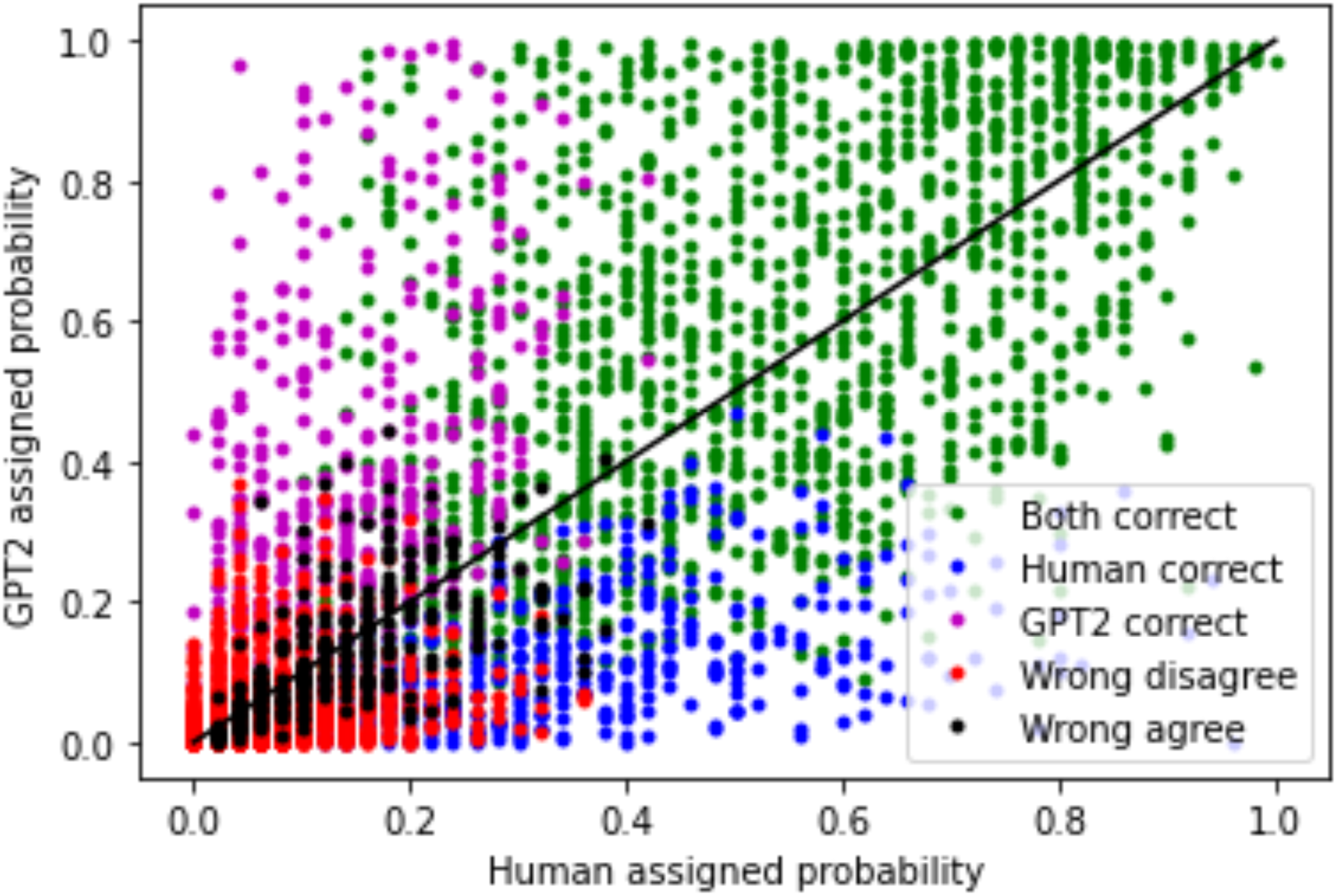
Comparing GPT-2 predictions and human predictions. **A)** Coloring the scatter plot according to GPT-2/human accuracy. GPT-2 and humans jointly predicted correctly 27.6% of the words (green). GPT-2 and humans jointly predicted incorrectly and disagreed on the next word for 48.8% of the words (red). GPT-2 and humans jointly predicted incorrectly and agreed on the next word for 5.9% of the words (black) 9.2% of the words humans predicted correctly were not correctly predicted by GPT-2 (blue). 8.4% of the words correctly predicted by GPT-2 were not correctly predicted by humans.

**Figure S3.**
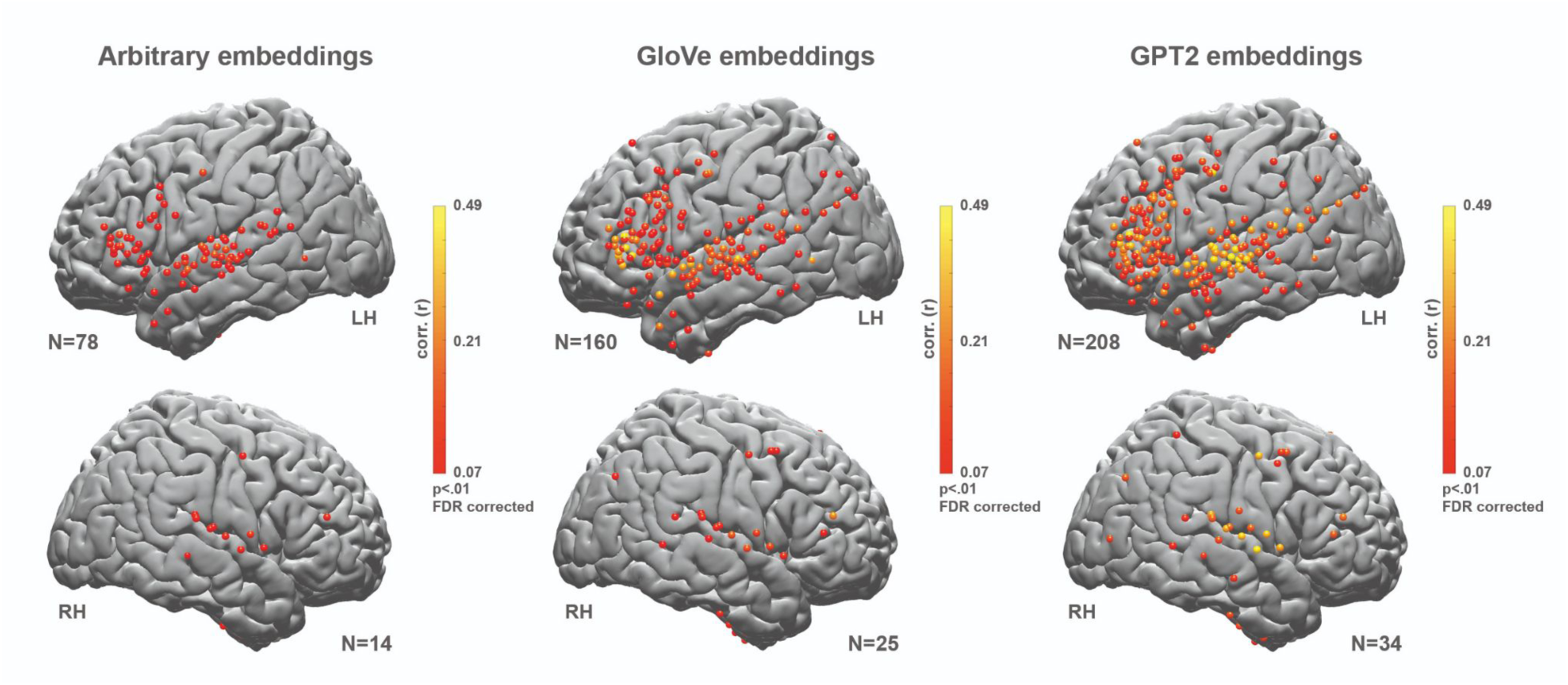
Left and right hemisphere encoding results show an advantage for contextual (GPT-2) embeddings over static (GloVe) and arbitrary embeddings. Right Hemisphere maps for correlation between. **A)** Predicted and actual word responses for the arbitrary embeddings (nonparametric permutation test; *q* < .01, FDR corrected). **B)** Correlation between predicted and actual word responses for the static (GloVe) embeddings. **C)** Correlation between predicted and actual word responses for the contextual (GPT-2) embeddings. Using contextual embeddings significantly improved the encoding model’s ability to predict the neural signals for unseen words across many electrodes. Given that we had fewer electrodes in the right hemisphere relative to the left hemisphere, this study is not set up to test differences in language lateralization across hemispheres.

**Figure S4.**
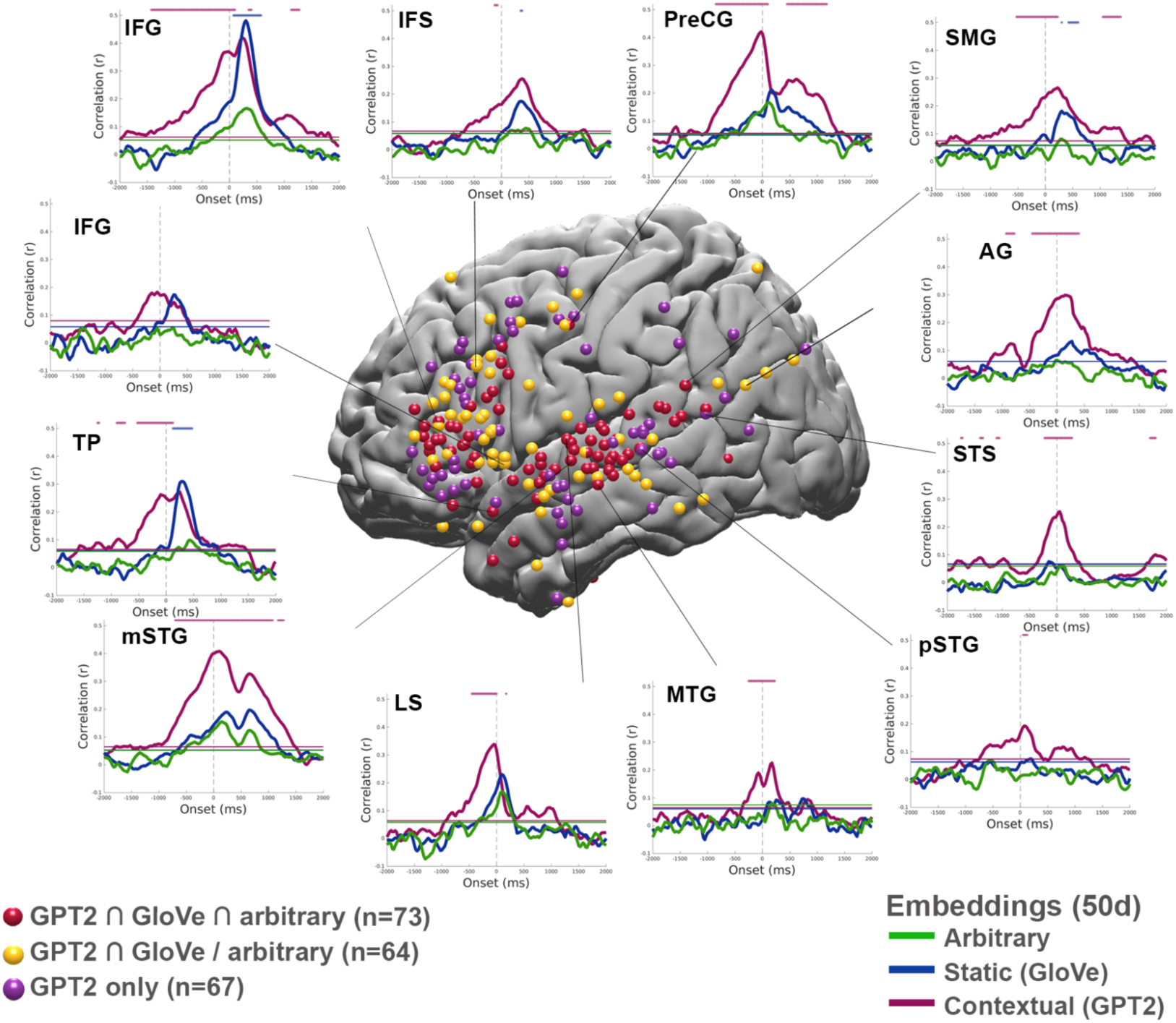
Contextual embedding significantly improves the modeling of neural signals. **A)** Map of the electrodes in the left hemisphere with significant encoding for 1) all three types of embeddings (GPT-2 n GloVe n arbitrary, red); 2) for static and contextual embeddings (GPT-2 n GloVe, but not arbitrary, yellow); 3) and contextual only (GPT-2, purple) embeddings. Note the three groups do not overlap. A sampling of encoding performance for selected individual electrodes across different brain areas: inferior frontal gyrus (IFG), temporal pole (TP), middle superior central gyrus (mSTG), superior temporal sulcus (STS), lateral sulcus (LS), middle temporal gyrus (MTG), posterior superior temporal gyrus (pSTG), angular gyrus (AG), post central gyrus (postCG), precentral gyrus (PreCG), and middle frontal sulcus (MFS). (Green -encoding for the arbitrary embeddings, blue - encoding for static (GloVe) embeddings; purple - encoding for contextual (GPT-2) embeddings).

**Figure S5.**
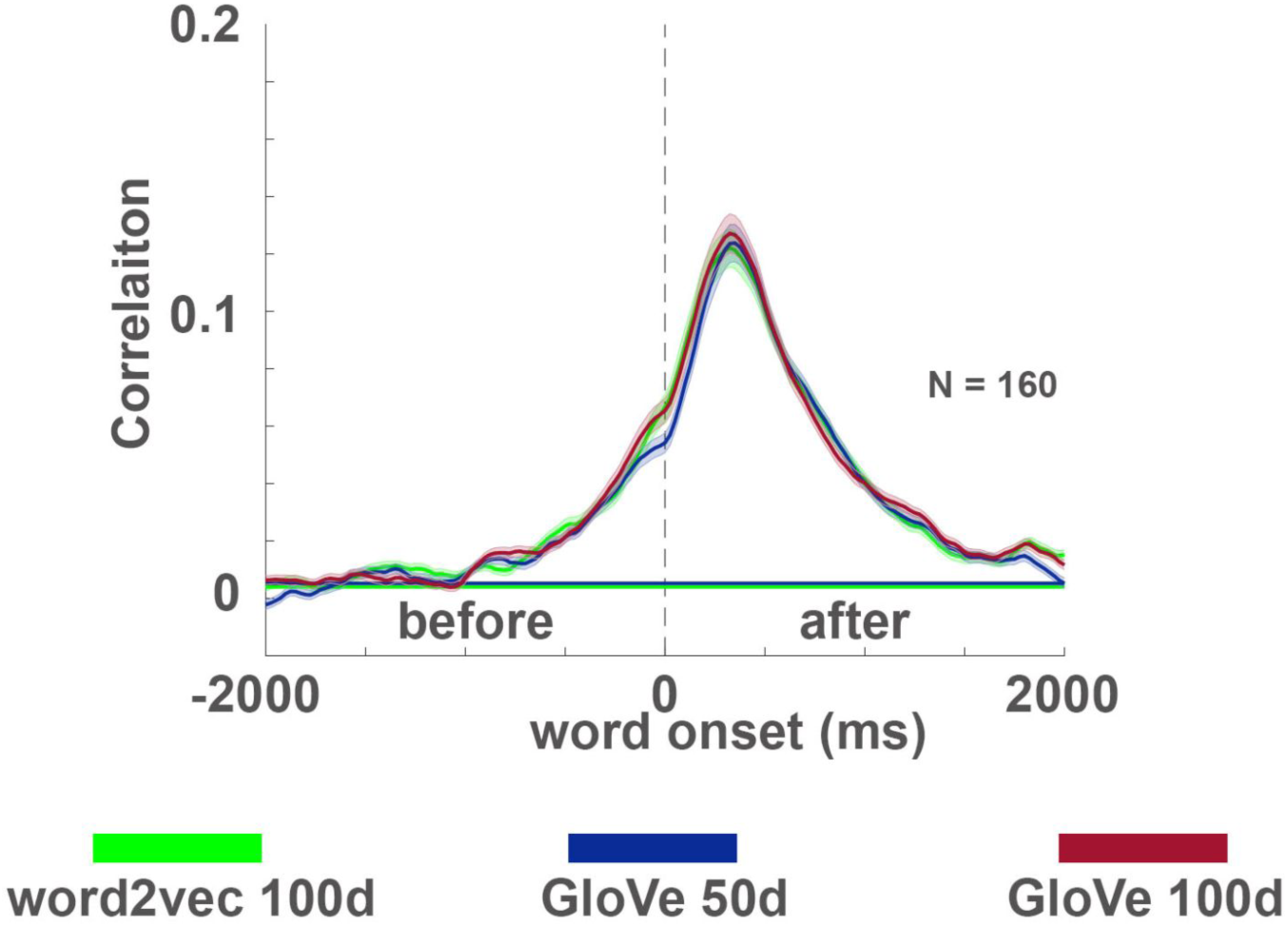
Comparison of GloVe- and word2vec-based static embeddings. The encoding procedure was repeated for two additional static embeddings using the electrodes that were found significant for GloVe-50 encoding on the left hemisphere (Fig. 3B). Each line indicates the encoding model performance averaged across electrodes for a given type of static embedding at lags from -2000 to 2000 ms relative to word onset. The error bars indicate the standard error of the mean across the electrodes at each lag. 100-dimensional word2vec and GloVe embeddings resulted in similar encoding results to the initial 50-dimensional GloVe embeddings. This suggests that results obtained with static embeddings are robust to the specific type of static embeddings used.

**Figure S6.**
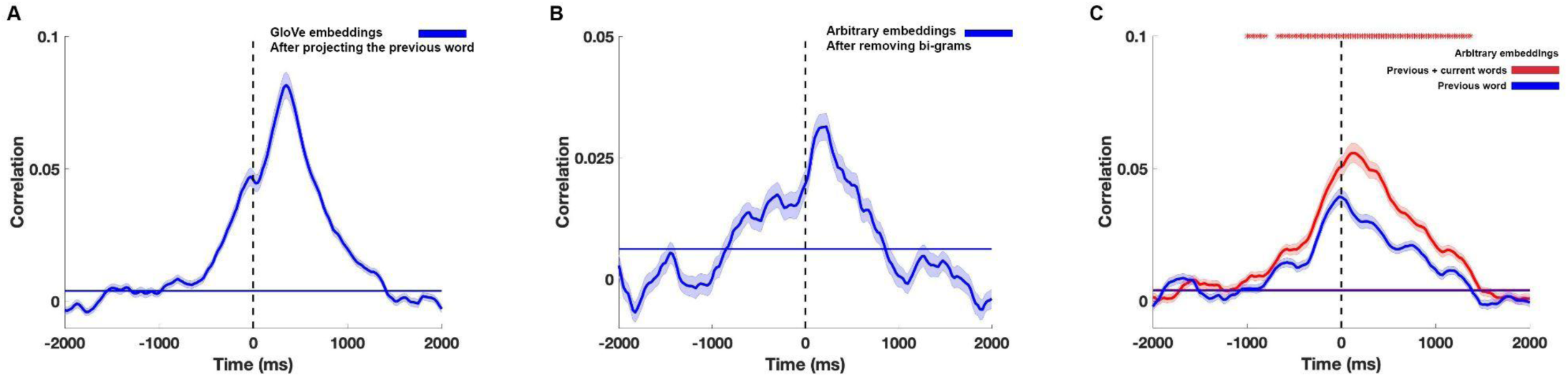
Controlling for correlations among adjacent GloVe embeddings. To ensure that the signal predicted before word-onset is not a result of a correlation among adjacent GloVe embeddings we ran the following additional control analyses: **A)** We projected (by inner product) and then subtracted the GloVe embedding of the previous word from each word and re-ran the encoding analysis. The analysis demonstrates that the significant encoding before word onset holds even after removing local contextual dependencies in the GloVe embedding of consecutive words. The horizontal line indicates the significance threshold calculated using a permutation test and FDR corrected for multiple comparisons (q<.01). **B)** We trained an encoding model using arbitrary embeddings on our dataset after removing the first word from all bi-grams that repeated more than once. The encoding before word onset remained significant after the removal of the bi-grams. The horizontal line indicates the significance threshold calculated using a permutation test and FDR corrected for multiple comparisons (q<.01). **C)** We compared an encoding model based on arbitrary embeddings using the previous word embedding (blue line), to an encoding model where we concatenated previous and current word embeddings (red line). Red asterisks mark significant differences using a permutation test and FDR correction (q<.01). The significant difference between these two models before word onset is another evidence that there is predictive information in the neural activity as to the upcoming word, above and beyond the contextual information embedded in the previous word. The horizontal line indicates the significance threshold calculated using permutation test and FDR corrected for multiple comparisons (q<.01).

**Figure S7.**
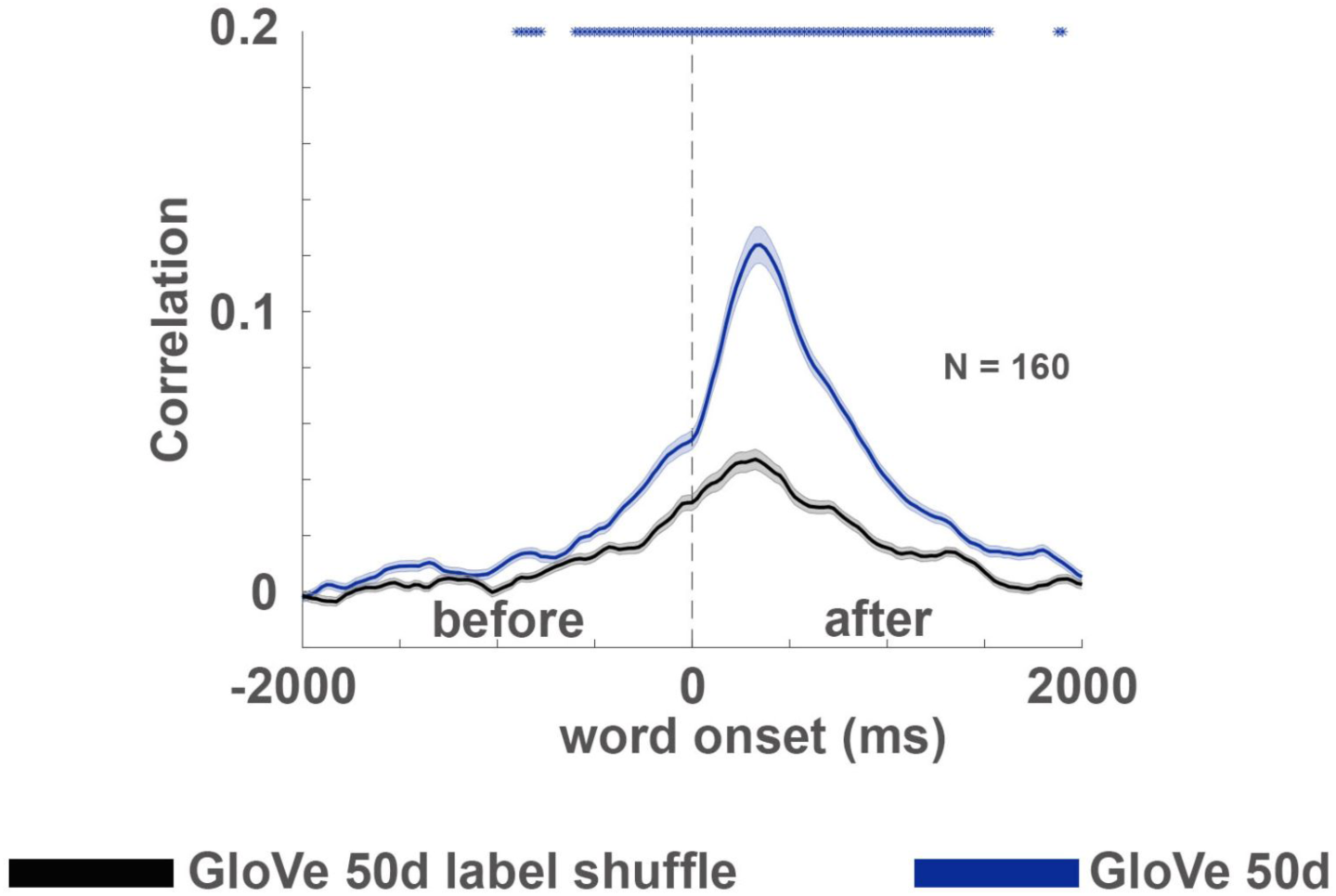
GloVe’s space embedding attributes. It can be argued that GloVe based encoding outperforms arbitrary-based encoding due to a general property of the space that GloVe embeddings induce (for example, they are closer / further away from each other). To control for this possible confound, we consistently mismatched the labels of the embeddings of GloVe and used the mismatched version for encoding. This means that each unique word was consistently matched with a specific vector that is actually an embedding of a different label (for example, matching each instance of the word ‘David’ with the embedding of the word ‘court’). This manipulation uses the same embedding space that GloVe uses and also induces a consistent mapping of words to embeddings (as in the arbitrary-based encoding). The matched GloVe (blue) outperformed the mismatched GloVe (black), supporting the claim that GloVe embedding carries information about word statistics that is useful for predicting the brain signal.

**Figure S8.**
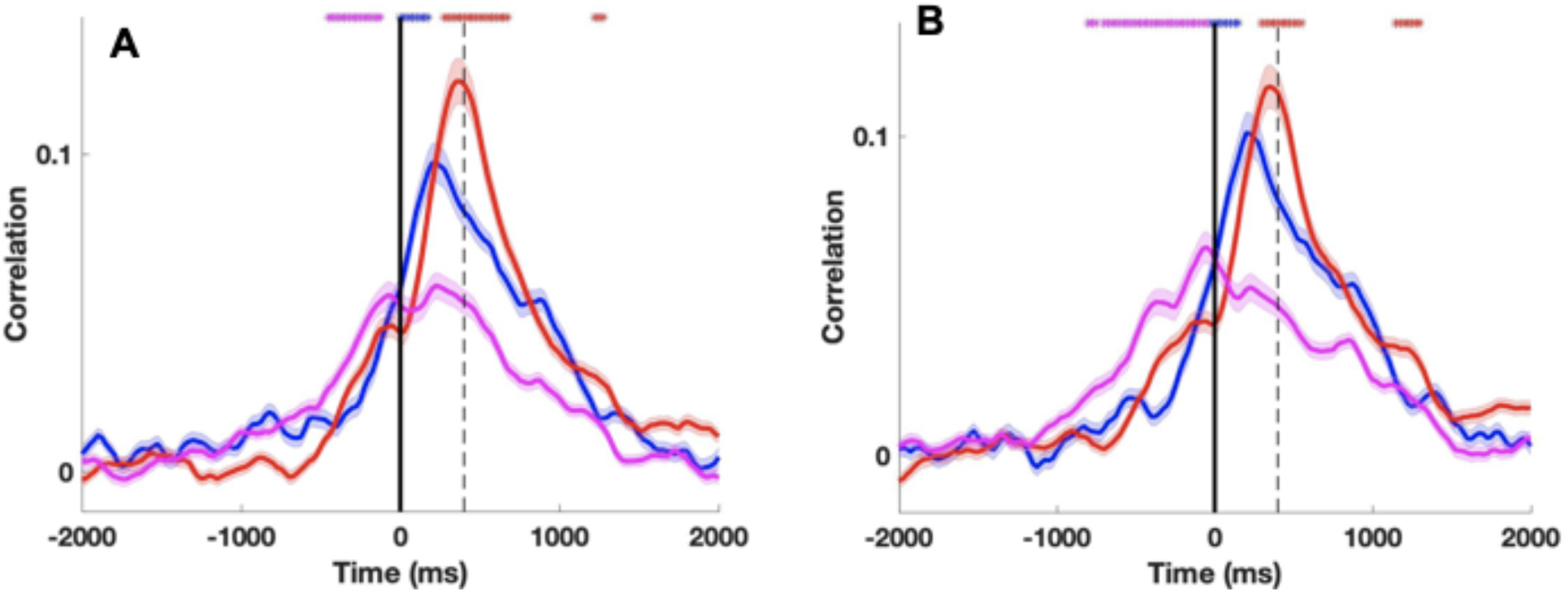
Encoding for correct / incorrect predictions. This is a variation of Fig. 4B where: A. We classify words as correctly predicted if they are the most predictable words by humans’ ratings. B. We classify words as correctly predicted if they are the most predictable by GPT-2 (instead of top-5).

**Figure S9.**
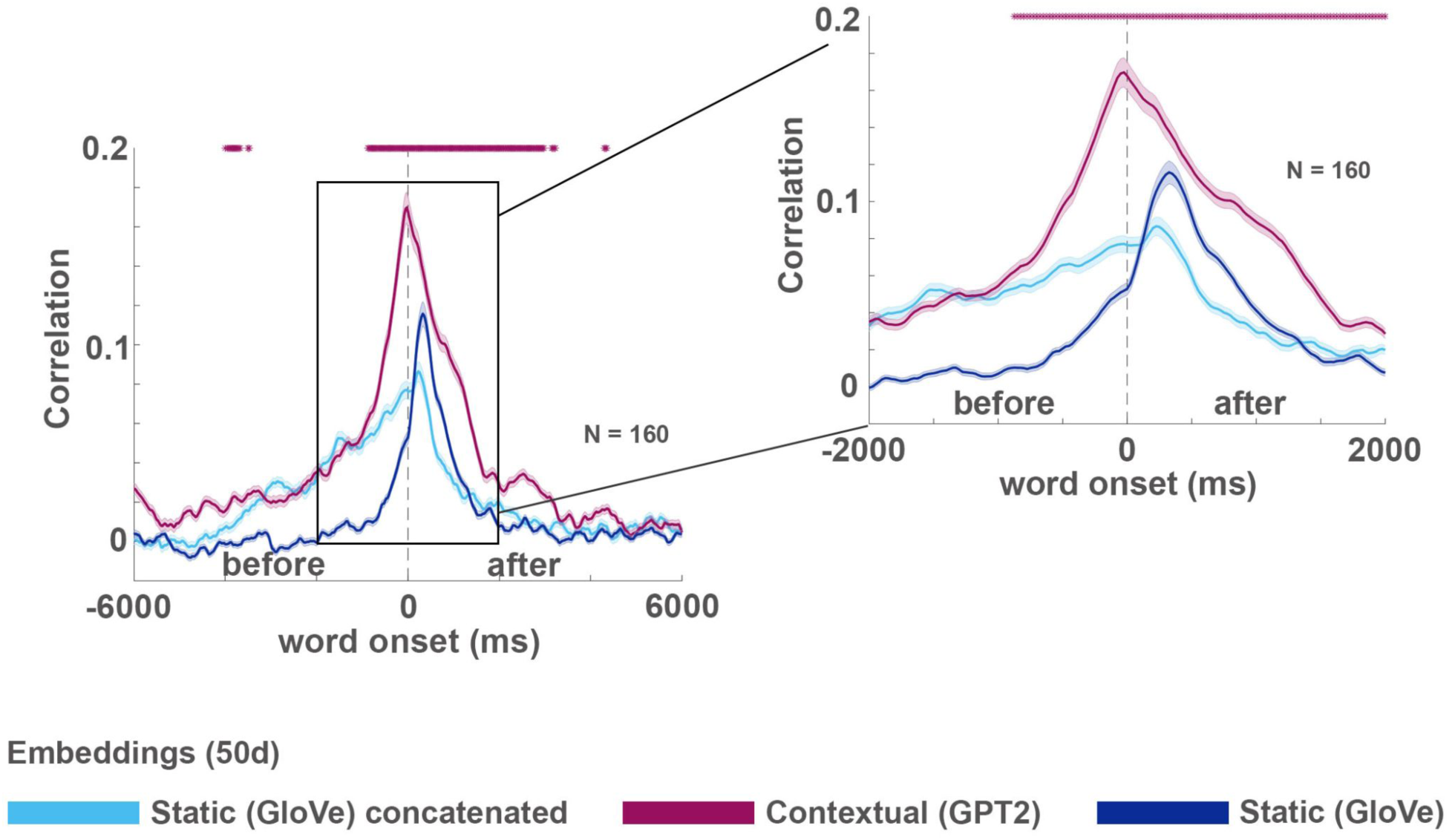
Comparison of GPT-2 and concatenation of static embeddings. The increased performance of GPT-2 based contextual embeddings encoding may be attributed to the fact that it consists of information about the previous words’ identity. To examine this possibility, we concatenated the GloVe embeddings of the 10 previous words and current word, and reduced their dimensionality to 50 features. GPT-2 based encoding outperformed mere concatenation before word onset, suggesting that GPT-2’s ability to compress the contextual information improves the ability to model the neural signals before word onset.

**Figure S10.**
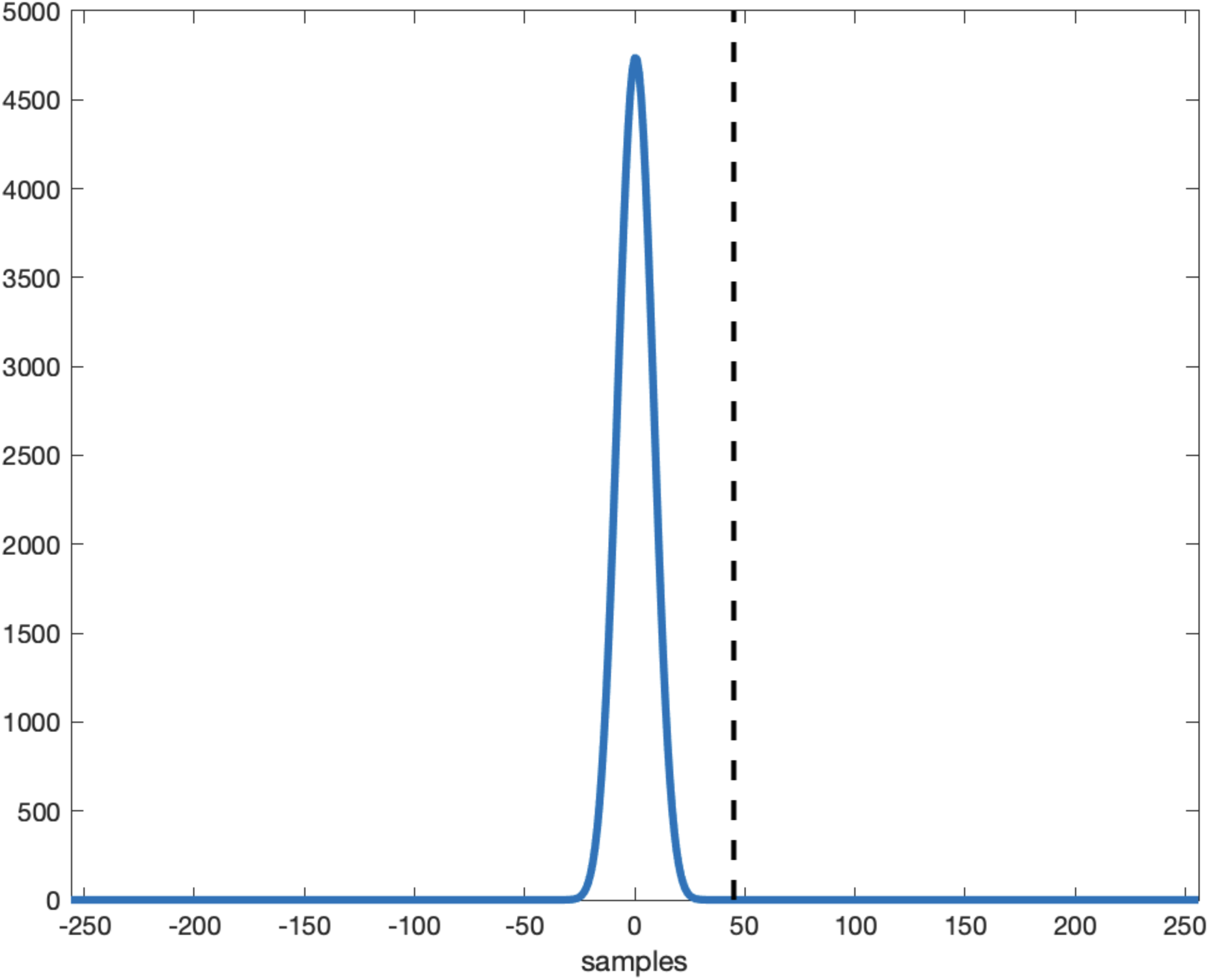
Preprocessing procedure applied to an impulse response. The plot demonstrates the temporal uncertainty introduced by the preprocessing procedure (especially by the wavelet and smoothing procedures). At sample 45 after onset (dashed line) the value is back to zero, considering the 512 HZ sampling rate this means that the leak from the future is bounded by 93 ms.

## Appendix I Decoding Model Details

*The size of the layer is dependent on the number of electrons included in the fold (5 folds over all). The number of electrodes ranged from 114-132, and the total number of parameters ranged from 219,670-228,790.

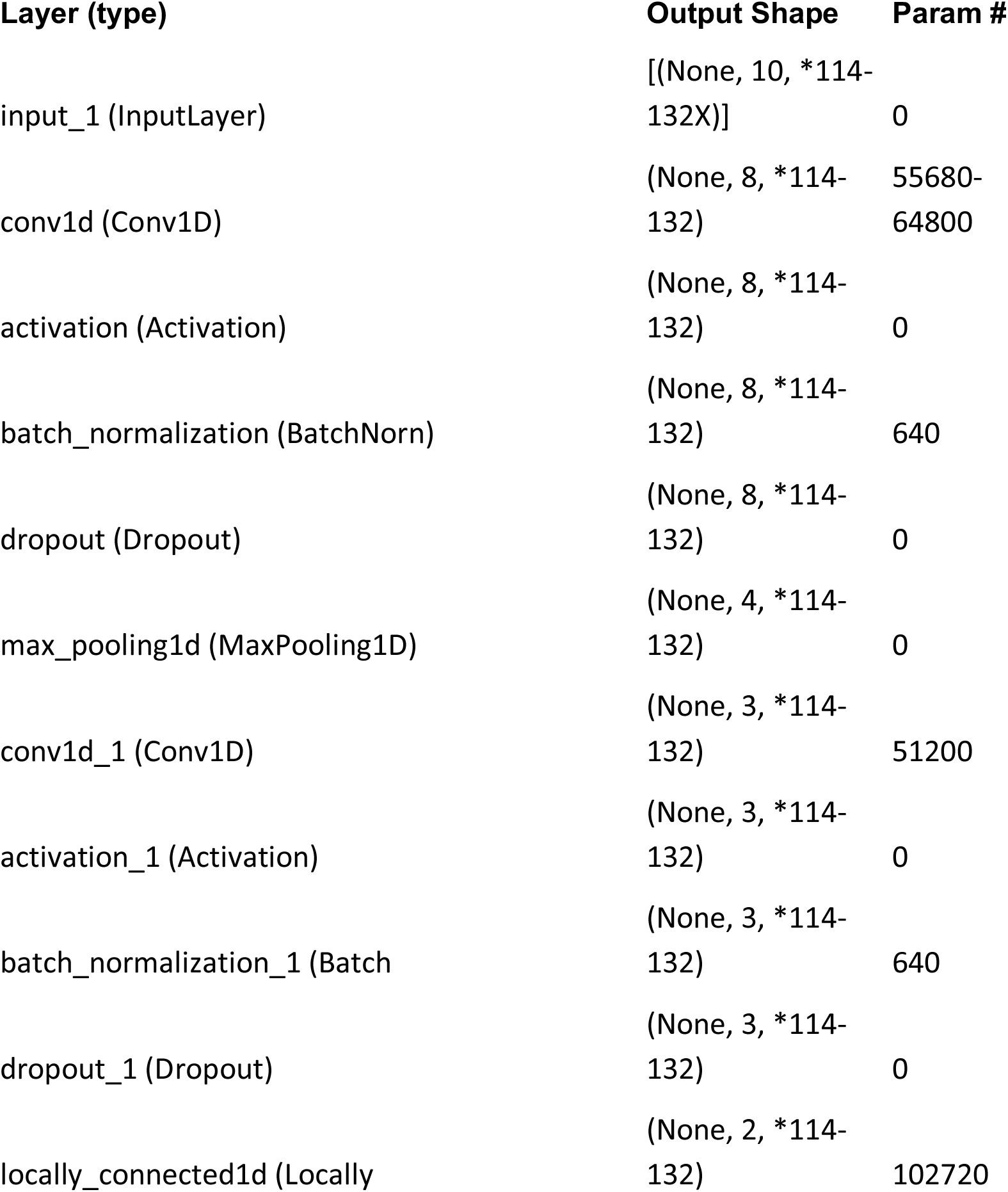
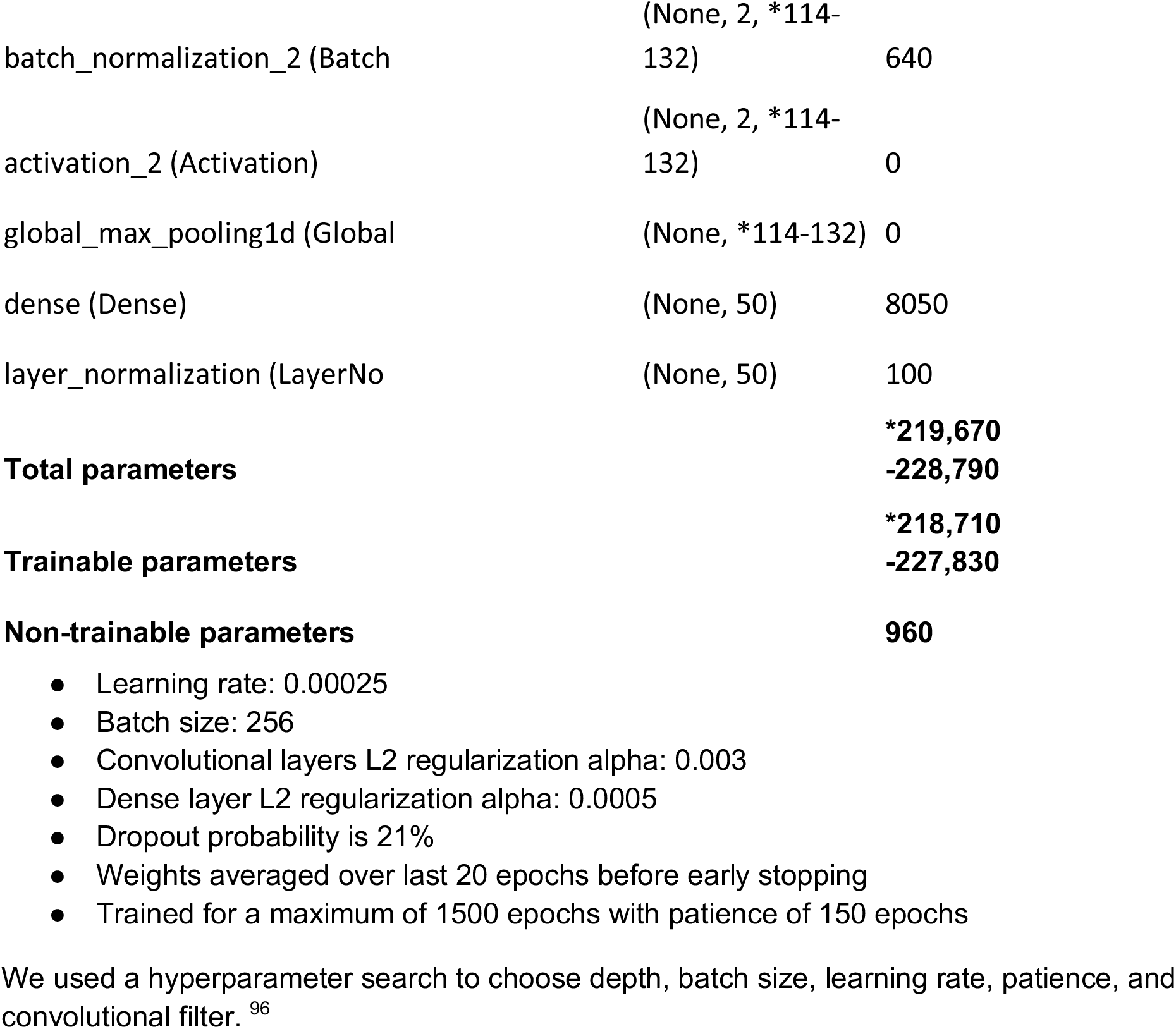

## Appendix II Word List

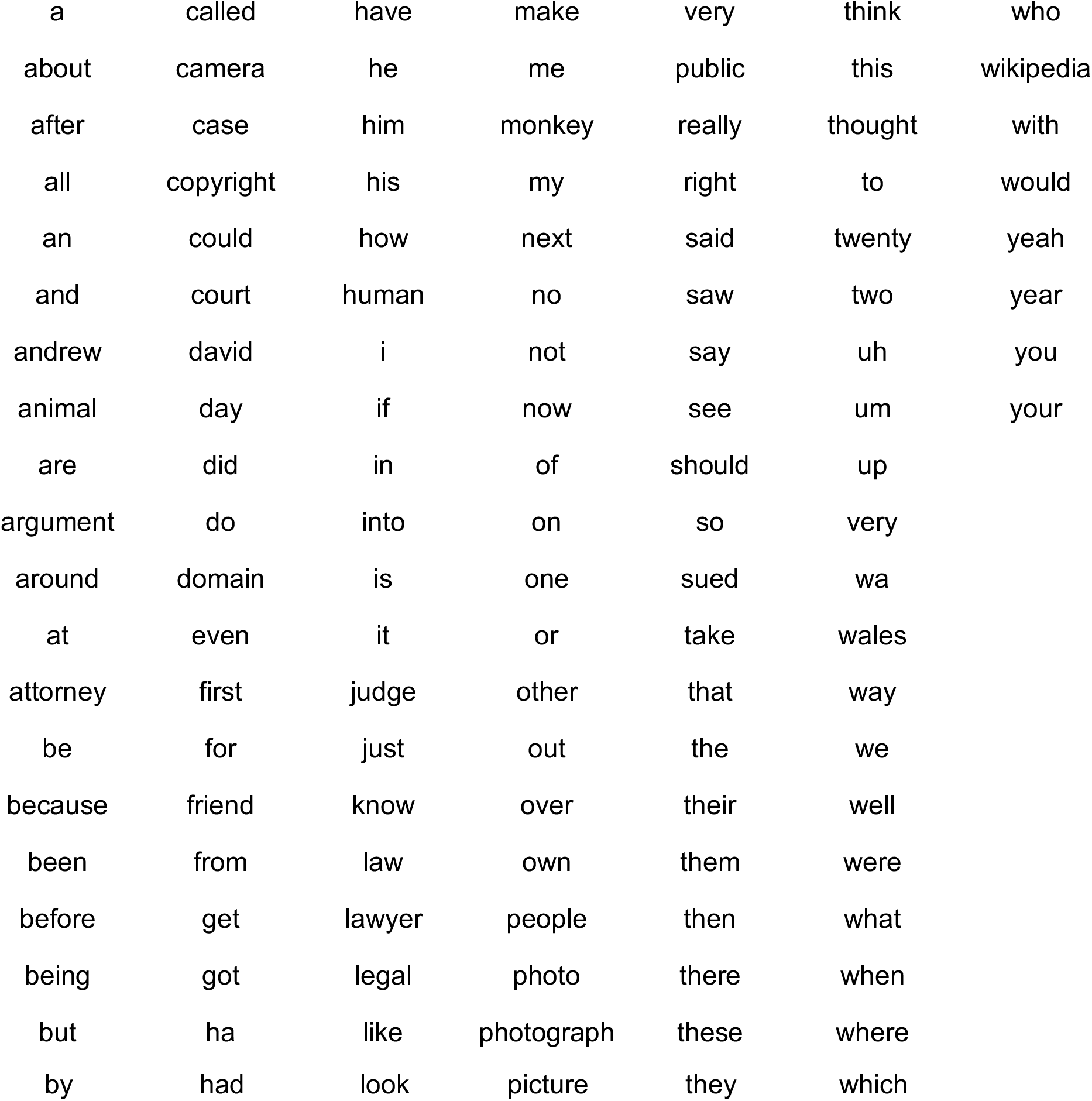

